# Molecular dependencies and genomic consequences of a global DNA damage tolerance defect

**DOI:** 10.1101/2023.10.11.561854

**Authors:** Daniel de Groot, Aldo Spanjaard, Ronak Shah, Maaike Kreft, Ben Morris, Cor Lieftink, Joyce J.I. Catsman, Shirley Ormel, Matilda Ayidah, Bas Pilzecker, Olimpia Alessandra Buoninfante, Paul C.M. van den Berk, Roderick L. Beijersbergen, Heinz Jacobs

## Abstract

DNA damage tolerance (DDT) enables replication to continue in the presence of fork stalling lesions. To determine the molecular and genomic impact of a global DDT defect, we studied *Pcna^K164R/-^;Rev1^-/-^* compound mutants. Double mutant (DM) cells displayed increased replication stress, hypersensitivity to genotoxic agents, replication speed, and repriming. A whole genome CRISPR-Cas9 screen revealed a strict reliance of DM cells on the CST complex, where CST promotes fork stability. Whole genome sequencing indicated that this DM DDT defect favors the generation of large, replication-stress inducible deletions of 0.4-4.0kbp, defined as type 3 deletions. Junction break sites of these deletions revealed preferential microhomology preferences of 1-2 base pairs, differing from the smaller type 1 and type 2 deletions. These differential characteristics suggest the existence of molecularly distinct deletion pathways. Type 3 deletions are abundant in human tumors, can dominate the deletion landscape and are associated with DNA damage response status and treatment modality. Our data highlight the essential contribution of the DDT system to genome maintenance and type 3 deletions as mutational signature of replication stress. The unique characteristics of type 3 deletions implicate the existence of a novel deletion pathway in mice and humans that is counteracted by DDT.

## INTRODUCTION

Genomes are subjected to a continuous barrage of genetic insults(1, 2). To preserve genetic information, genome maintenance is essential for life. A versatile DNA damage response (DDR) network warrants that most genotoxic lesions are sensed and restored effectively(3, 4). Defects in DNA repair pathways can result in accelerated ageing, increased tumorigenesis, sensitivity to DNA damage and various developmental defects(2, 3, 5–8). As DNA lesions usually arise in one of the two complementary strands, the repair synthesis of excision-based repair pathways strongly relies on the integrity of the complementary DNA strand. During replication, however, strand separation prohibits canonical excision repair of DNA lesions that persisted into S phase. These lesions can stall replicative DNA polymerases, which may result in a collapse of the replication fork and generation of more mutagenic secondary lesions such as DNA double strand breaks (DSBs)(9). DSBs arising in S-phase are repaired in a relatively error-free manner by homology directed DSB repair (HR) which uses the intact template of the sister chromatid to bypass the lesion. However, alternative, more error-prone DSB repair pathways such as canonical non-homologous end joining (NHEJ) and alternative end joining (A-EJ) are active as well. These are associated with a broad spectrum of mutations, ranging from subtle InDels to structural variances such as deletions, chromosome rearrangements, and translocations(10, 11).

To bypass replication blocking lesions and simultaneously prevent recombinogenic DSBs, cells employ several DNA damage tolerance (DDT) pathways that enable replication to continue opposite DNA lesions, thereby stabilizing the replisome, thought to come at the expense of accuracy(10, 12–14). In addition, DDT enables replicative bypass of unhooked interstrand crosslink (ICL). In this way, DDT constitutes an integral, intermediate step in ICL repair pathways, including Fanconi anemia repair(2, 15, 16). Furthermore, DDT can contribute to nucleotide excision repair (NER)(17). Four distinct modes of DDT are distinguished: (i) translesion synthesis (TLS) opposite the lesion, (ii) template switching (TS), which takes advantage of the intact template of the sister chromatid avoiding the lesion, (iii) fork reversal (FR), during which the replication fork is remodeled into four-way junctions to avoid the lesion, and (iv) repriming behind the damaged template followed by gap filling TLS (‘post-replicative’ TLS) or by homology directed repair of the remaining gap (‘post replicative’ TS)(18).TLS utilizes specialized DNA damage tolerant Y-family DNA polymerases η (POLH), ⍳ (POLI), ϰ (POLK) and REV1 and the B-family polymerase ζ (POLZ), which allows replication to continue directly opposite specific lesions(12, 19)(19, 20). TLS polymerase recruitment is facilitated by damage-induced *mono*-ubiquitination of PCNA (PCNA-mUb) on lysine residue 164(21, 22). TS and FR depend on damage-induced *poly*-ubiquitination of PCNA (PCNA-Ub^n^) at lysine residue 164 and avoid the lesion by continuing DNA synthesis on the newly synthesized intact template of the sister chromatid(23)(23). TLS polymerase REV1 has been implicated in the recruitment and activation of other Y-family TLS polymerases independent of PCNA-Ub through its C-terminal domain(24–27)(24–27).

TLS is considered to be the predominant mode of DDT in mammals(28). TLS polymerases -except REV1-feature a capacious active site that can incorporate bulky nucleotides and non-Watson-Crick base pairs(29–31). Moreover, in contrast to replicative polymerases, TLS polymerases are proof-read incompetent as they lack the 3’-5’ exonuclease domain to remove ‘misincorporated’ bases. Consequently, as shown in gapped plasmid assays, replication opposite non-instructive lesions can be highly error prone(20, 32–34). Based on these *in vitro* observations TLS, as predominant arm in mammalian DDT, is in general considered mutagenic. In contrast, TLS can also be error-free, as is the case for POLH, which can tolerate specific UV lesions(35). Xeroderma Pigementosum-Variant (XP-V) patients that lack POLH activity have an increased risk of developing skin cancer upon UV exposure(5). DDT is observed in all domains of life; is an important building block of the mammalian DNA damage response network and essential for mammalian life(36, 37). The sum of these insights suggests an important contribution of DDT to genomic integrity in the context of genotoxic lesions and fork-stalling structural impediments in the DNA template.

Our previous studies documented an essential, non-redundant contribution of PCNA-Ub and REV1 controlled DDT for mammalian development, stem cell maintenance, and drug responsiveness(36, 38). We here defined the molecular and genomic impact of a compound DDT defect. To accomplish this goal, we generated lymphoma cell lines that were deficient for both PCNA-Ub and REV1. These mutations were found associated with increased sensitivity to most replication blocking agents, replication stress, and replication fork speed. The latter related to increased single strand gaps, indicating increased repriming activity as a remaining alternative to bypass lesions to warrant fork stability. Furthermore, to define molecular dependencies and backup mechanisms of cells suffering from a global DDT defect, we performed an unbiased genome wide CRISPR-Cas9 drop out screen. This screen identified the CST complex, comprising CDC13/CTC1, STN1, and TEN1, as essential for cell survival and cell cycle progression of DM cells. CST is involved in telomere capping and regulates telomerase and DNA polymerase alpha-primase (polα-primase) access to telomeres(39). Apart from its telomere function, the CST complex mediates DNA end protection by blocking MRE11 mediated degradation of nascent DNA strands, thereby protecting stalled replication forks(39–41). Recently, CST has also been found to directly control fork progression through modulation of DDR activity(42), to counteract toxicity of oxidative damage(43), and to be required for survival in yeast deficient for Rad6(44). Though a role of CST in repriming has been proposed(45), our DNA fiber assays rather support a function in fork stabilization. Surprisingly, further genomic characterization of *Pcna^K164R/-^*;*Rev1*^-/-^mutation, revealed that the accumulation of small-scale mutations was not grossly altered, as shown by whole genome sequencing (WGS). Instead, our analyses revealed a multimodal size distribution of genomic deletions, each of which was characterized by a preferential 1-3 bp micro-homology pattern at the break/junction sites. Notably, the DDT defect favored a specific accumulation of UV damage inducible, large type 3 deletions ranging between 0.4-4.0 kbp in size, likely as a consequence of replication stress. To extrapolate the impact and relevance of our findings to the human system, we took advantage of the WGS database of human tumors as provided by the Hartwig Medical Foundation (HMF). Remarkably, deletions of similar size were found abundant in human cancers. These deletions appear to increase in the presence of DDR defects and decrease upon amplification of defined DDR genes and appear affected by the treatment modality. The sum of our insights highlights the critical contribution of the DDT system in genome maintenance, both in human and mice. A multimodal size distribution of deletions and selective micro-homology preferences implicate distinct repair activities underlying their generation. We propose type 3 deletions as a replication stress associated and damage-inducible mutational feature, which suggests the existence of a specific deletion pathway that is counteracted by DDT.

## MATERIALS AND METHODS

### Derivation of isogenic DDT proficient and deficient mouse thymic lymphoma cell lines

To obtain WT and *Pcna^K164R/-^*;*Rev1*^-/-^ lymphoma cell lines, 10 x 10^6^ lymphoma cells from a p53-KO, *Pcna^K164R/loxP^* mouse(46) were nucleofected. A nucleofector II-b device (Amaxa, Lonza), program A-30 with 10 µg of plasmid DNA and 1 µg of eGFP control plasmid (Amaxa, Lonza) was used. Subsequently, GFP or mCherry positive cells were sorted with MoFlo Astrios or FACSAria IIu (BD Biosciences) cell sorters. Clones were obtained via limiting dilution. To obtain DM lymphoma cells, *Pcna^K164R/-^* mutant cells were transfected with the indicated guides against *Rev1*.

### Generation of mouse embryonic fibroblasts

Primary MEFs were isolated from E14.5 embryos intercrosses of *Pcna^K164R/+^*;*Rev1*^+/-^ and *Pcna^K164R/+^*;*Rev1*^-/-^ mice. Lentivirally transduction with p53 shRNA(47) was performed in order to immortalize the MEFs.

### gRNA design to target murine *Pcna* and *Rev1*

Benchling was used to design guides targeting two exons or introns of each gene. For *Pcna*, the following guides were used: 5’-TAGTAAGGGGGCGTCCAGTT-3’ and 5’-GAATTTTGGACATGCTAGGG-3’. For *Rev1*, the same guides were chosen that had been used to generate the *Rev1^-/-^* mouse: 5’-AGAAATCTAATGATGTTGCATGG-3’, and 5’-TGAAGCACTGATTGACGTCACGG-3’. These were cloned into pX330 (one guide per plasmid) or a modified form of pX330, named pX333-mCherry, which contained sites for two guides. The protocol from Ran *et al.* was used to clone the guides in pX330(48).

### Isolation of genomic DNA and PCR

To obtain DNA for mouse-genotyping, and to screen for the presence of mutations in *Pcna*- or *Rev1* in lymphoma cells, cells or mouse toes were lysed with ProtK (10 mg/mL). Mutations were screened for using PCR (see Supplementary table 1 for primers and table 2 for PCR settings).

### Proliferation assay MEFs

Viable primary MEFs or p53kd MEFs were seeded at the concentration of 300 cells per well in 96 well plates. The cells were allowed to grow for 8 days or 10 days in IncuCyte® (Essenbioscience) under standard tissue culture conditions. Images were taken every 4 hours and results analyzed in Excel (Microsoft) as percentage of confluence, that represents the percentage of the image area occupied by objects.

### Survival assay, synergy experiments, and high-throughput compound screen

For survival assays in the lymphoma cell lines, cells were seeded at low confluency using the Multidrop Combi (Thermo Fisher Scientific), into 384-well plates (Greiner). After 24 h, the respective compounds were added using a tecan d300e compound printer (HP), including 3–5 replicates per dosage. Positive (1 μM phenylarsine oxide) and negative (0, 1% DMSO) controls were added to each assay plate. Viability readout was performed after 72 h, using the CellTiter-Blue assay (G8081/2, Promega) following the protocol of the manufacturer using an EnVision multimode plate reader (PerkinElmer). The CTB data was normalized per plate using the normalized percentage inhibition (NPI) method(49). NPI sets the mean of the positive control value to 0 and mean of the negative control to 1. Synergy calculations were performed in R using the Chou-Talalay Combination index method cytometry. UV survival assays were performed by seeding 500, 000 cells in 24 well plates and treated with UV-C. After 72 hours, viability was assessed by flow cytometry. For the high-throughput drug screen in lymphoma cells, the following procedure was followed: Using the Multidrop Combi (Thermo Fisher Scientific), untreated p53-KO lymphoma cells were seeded into 384 well plates at low confluency. After 24 hours, the collection of compound libraries available at the NKI (Selleck GPCR, Kinase, Apoptosis, Phospatase, Epigenetic, LOPAC, and NCI oncology) was added. This library was stored and handled as recommended by the manufacturer. Compounds from the master plate were diluted in daughter plates containing complete RPMI 1640 medium, using the microlab star liquid handling workstation (Hamilton). From the daughter plates, the diluted compounds were transferred into 384 well assay plates, in triplicate, with final concentrations of 1 μM, and 5 μM. In addition, Positive (1 μM Phenylarsine oxide) and negative (0.1% DMSO) controls were added alternately to wells in column 2 and 23 of each assay plate. After 3 days, viability was measured using CellTiter-Blue assay (G8081/2, Promega) following the protocol of the manufacturer. The CTB data was normalized per plate using the normalized percentage inhibition (NPI) method. NPI sets the mean of the positive control value to 0 and mean of the negative control to 1. Using the replicate values of both conditions a two-sided *t*-test was performed. Afterwards the *p-*values were corrected for multiple testing using the Benjamini-Hochberg method. All calculations were done in R.

### Measure of replication stress by yH2AX

Lymphomas were plated in 24 well plates with 500.000 cells per ml in 0.5 ml medium. After 15 minutes cells were exposed to 20 J/m^2^ UV-C and given 0.5ml of fresh medium. To distinguish dead cells, cells were stained with LIVE/DEAD Fixable Yellow Dead Cell Stain Kit (Thermo Fisher) for 20 minutes. After this, cells were incubated at 37°C for 4 hours after which they were fixed and permeabilized with the Transcription Factor Buffer Set from BD. γH2AX was detected with PerCP/Cyanine5.5 anti-H2A.X-Phosphorylated (Ser139, clone 2F3, Biolegend) antibody (1:500). Finally, cells were resuspended in PBS with 7 μg/ml DAPI and measured on an Attune (Thermo Fisher). Data were analyzed using FlowJo software version 10.7.0.

### EdU incorporation assay

Cells were labeled with 1:1000, 10 mM EdU in regular medium for 20 minutes after which cell were washed, resuspended in PBS and fixed in 100% ethanol. The Click-iT Plus EdU Flow Cytometry Kit AF488 (Thermo Fisher Scientific) protocol was followed. Cells were suspended in PBS containing 7 μg/ml DAPI to visualize DNA content. Finally, samples were measured with an Attune (Thermo Fisher) and analyses were performed using FlowJo version 10.7.0.

### Whole genome dropout screen

The Mouse Improved Genome-wide Knockout CRISPR Library v2 (Addgene #67988) collection sgRNAs in lentiviral vectors targeting 18, 424 mouse genes, with four to five individual sgRNAs for each gene, was used to infect WT and PcnaK164RRev1 -/- lymphoma with a 100-fold coverage. Lymphoma were harvested at T = 0 after a three-day period in which cultures underwent selection with 2µg/ml puromycin to select for cells that had been transduced with the library. Cells were subsequently cultured for ≈10 cell divisions. After 10 doublings, gDNA was isolated using the DNAeasy blood & tissue kit (QIAGEN) following the manufacturer’s protocol. For each line, one replicate was taken at T=0, and three replicates were taken at T=10. A capture protocol was performed to enrich for sgRNA sequences. gDNA was digested overnight at 37°C with NdeI and XhoI. The following day, samples were heated for 10 minutes at 100°C, and subsequently snap frozen. After light thawing, biotinylated capture oligos were added to enrich for sgRNA-barcodes: NdeI-Fwd: 5’-TGCTTACCGTAACTTGAAAGTATTTCGATTTCTTGGCTTTATATATCTTG-3’ and XhoI-Rev: 5’-GATCTAGATGGATGCAGGTCAAAGGCCCGGACATGAGGAAGAGGAGAA-3’. Samples were hybridized overnight at 60°C. Biotinylated primers were captured using Streptavidin T1 dynabeads for 2 hours at room temperature. After multiple washes, beads were used as input material for subsequent PCRs: sgRNAs were retrieved using Phusion High-Fidelity DNA Polymerase (Thermo Fisher Scientific) by a 2-step PCR protocol. First, biotinylated capture oligos were amplified in the first round of PCR amplification with 10 μl 5× GC buffer, 1 μl 10 mM dNTP, 2.5 μl 10 μM PCR 1 indexed forward primer (Table S2, primers TRC_PCR1_Fwd1-8, which contain the barcode to label each sample), 2.5 μl 10 μM PCR 1 indexed reverse primer (Table S2, primer Yusa_Rv_1st PCR), 1.5 μl DMSO, 1 U Phusion Polymerase, and H2O added up to 50 μl and run at (1) 98°C, 30 s; (2) 98°C, 30 s; (3) 60°C, 30 s; (4) 72°C, 1 min (steps 2, 3, and 4 for 20 times) and 72°C, 5 min. The product of PCR1 was used to setup the second PCR reaction similar to PCR1. In PCR2, primers containing P5 and P7 sequences were used. The abundance of the guide RNAs in the T = 0 and T10 replicates was determined by Illumina Next Generation Sequencing. For the analysis of the screen, a differential analysis on the sgRNA level with DESeq2 was performed63. As hits, we selected genes for which at least two sgRNAs had a log2 fold change smaller than −1 and a FDR ≤ 0.1.

### Sample preparation and immunoblotting

To validate the genetic ablation of STN1, lymphoma cells were lysed in RIPA buffer (25 mM Tris HCl (pH 7.6), 150 mM NaCl, 1% NP-40, 1% sodium deoxycholate, and 0.1% SDS) containing 1x protease inhibitor cocktail (Roche) for 30 minutes on ice. Lysates were sonicated for 15 minutes using a BioRuptor (30 seconds on, 30 seconds off, maximum power, at 4°C). Samples were spun at 20, 000 g for 10 minutes, and the protein concentration was measured either via the Bradford assay, or via a BCA assay. Samples were run on a 4-12% Bis-Tris gel with MES running buffer on ice, at 150-200 V for 1.5-2 hours, until the loading band ran off the gel. Following, an overnight wet transfer was used. After staining with Ponceau-S, samples were blocked for 1 hour using PBST containing 5% skim-milk powder, followed by incubation with primary antibodies *Obfc1* Ab (sc-376450 1:250) for 3 hours at room temperature in TBS-T 1% milk. Blots were washed 3 times for 10 minutes with PBST, followed by a secondary staining (IRDye800CW goat anti-mouse, 1:5000 in PBST) for 1 hour at room temperature. The membrane was washed 3 times with PBST for 10 minutes, after which the membrane was imaged on an Oddyssey scanner (LiCor). For loading control GAPDH (14C10) was used (1:5000) as primary and IRDye680RD goat anti-rabbit (1:10000) as secondary.

### DNA fiber and S1 nuclease assay

In a 24 well plate 300.000 cells were plated in 250 μl medium. Prior to UV-C exposure (20 J/m^2^), lymphomas were incubated in medium containing 25 μM 5-Chloro-2-deoxyuridine (CldU) for 20 min at 37°C. After UV-C exposure, medium containing 500 μM 5-Iodo-2-deoxyuridine (IdU) was added, resulting in a final concentration of 250 μM IdU and 12.5 M CldU. After 60 min at 37°C, 10ml of 4°C PBS was added and cells were spun down at 1200rpm for 5 minutes and resuspended in 1ml PBS with 1% FCS. 2 µL of a suspension of 3 × 10^5^ lymphomas per mL was spotted onto a microscope slide, incubated for 5 min and lysed with 7 µL lysis buffer (200 mM Tris-HCl pH7.4, 50 mM EDTA, 0.5% SDS) for 3 min. Slides were tilted to 15°C to allow the DNA to run down the slide. Next, the slides were air dried and subsequently mixed in methanol-acetic acid (3:1). After rehydration, fixed DNA fibers were denatured in 2.5 M HCl for 75 min. Incorporation of CldU was detected using rat-anti-BrdU antibodies (1:500; BU1/75, AbD Serotec) and Alexa fluor-555-labelled goat-anti-rat antibodies (1:500; Molecular Probes), whereas incorporated IdU was detected using mouse-anti-BrdU antibodies (1:750; Clone B44, BD) and Alexa fluor-488-labelled goat-anti-mouse antibodies (1:500; Molecular Probes). Finally, slides were mounted in Fluoro-Gel (Electron Microscopy Sciences). Microscopy was performed using a fluorescent microscope (Zeiss). For the S1 nuclease assay cells were permeabilized with NuEX buffer after the staining and spinning. Next cells were washed with S1 nuclease buffer and spun down at 7000 rpm for 5 minutes. 20 U/ml S1 in S1 buffer was added for 30 min at 37°C, after which cells where washed and resuspended in 1 ml PBS with 1% FCS. Following steps were performed as normally.

### Whole genome sequencing, mutation calling, and analysis

WT and DM lymphoma cells were either irradiated one time with UV-C (0.4 J/m^2^) or left untreated. After, cells were left to recover for three days in regular medium, clones were obtained via limiting dilution. DNA was isolated using the Isolate II Genomic DNA kit (Bioline). Library preparation and whole genome sequencing was performed at the Hartwig Medical Foundation using the HiSeqX system generating 2 × 150 base read pairs. Sequencing quality control was done with FASTQC, followed by aligning by BWA(50) using the mm10 genome as reference. Finally, Mark Duplicates by Picard was used to remove duplicated reads. For structural variant calling GRIDSS(51) (version 2.1.0-0)was used and for small scale mutations GATK Haplotypecaller(52) (version 4.2.3.0) was used. Downstream analysis, quantification and visualization was done in R (version 4.2.0). MutationalPatterns (version 3.6.0) package was used for generating the base substitution signatures plots.

### Analysis of human whole genome sequencing data

The Hartwig Medical Foundation(53) database containing >4000 whole genome sequenced human tumors was used for these analyses. For downstream quantification and visualization R was used.

### Statistical analysis

To assess the statistical significance of our data we used the *t-*test, multiple unpaired student *t*-test with two-stage step-up correction (Benjamini, Krieger, and Yekutieli), one-way ANOVA, or Mann–Whitney *U* test (*P < 0.05, **P < 0.01, *** P < 0.001, **** P < 0.0001) by GraphPad Prism 6 or 9. No statistical methods were used to predetermine sample size.

## RESULTS

### DDT-deficient cells lacking PCNA-Ub and REV1

Previous work in our lab established a non-epistatic and embryonic lethal interaction of a *Rev1^-/-^* and the *Pcna^K164R/K164R^* mutation(7, 36). In contrast to single mutants, a compound mutant was found to be synthetic lethal in primary MEFs (Supplementary Fig. 1A). Ablation of p53 rescued this synthetic lethality (Supplementary Fig. 1A). Based on this knowledge, we made a thymic lymphoma cell line that was established from a *p53^-/-^;Pcna^K164R/loxP^* mouse, as described previously(46). Using this as a founder cell line we established genetically defined isogenic *Pcna*^wt/-^;*Rev1*^wt/wt^ (WT) and a double mutant *Pcna^K164R/-^, Rev1^-/-^* (DM) lymphoma cell lines. This provided a useful tumor model to study the relevance of DDT in DNA damage response and mutagenesis. As expected, a compound mutant DDT defect led to a profound increase in the sensitivity to various genotoxic agents (Figure 1A). The DM cells were hypersensitive to UV-C induced lesions, cisplatin, and the alkylating agent methy-methane sulfonate (MMS). In line with these results, an unbiased high throughput compound screen revealed many more compounds to which specifically DM lymphomas were highly sensitive. These either directly damage DNA (e.g. carboplatin, RITA NSC65287, and melphalan), or alter DNA metabolism (e.g. 6-thioguanine), or act as cell cycle inhibitors (CB1954) (Supplementary Fig. 2A-D). Intriguingly, one of the only compounds DM cells appeared more resistant to then WT cells was 5-Fluorouracil (Supplementary Fig. 2C, 2D), a finding consistent with a previous study(54).

**Figure 1.**
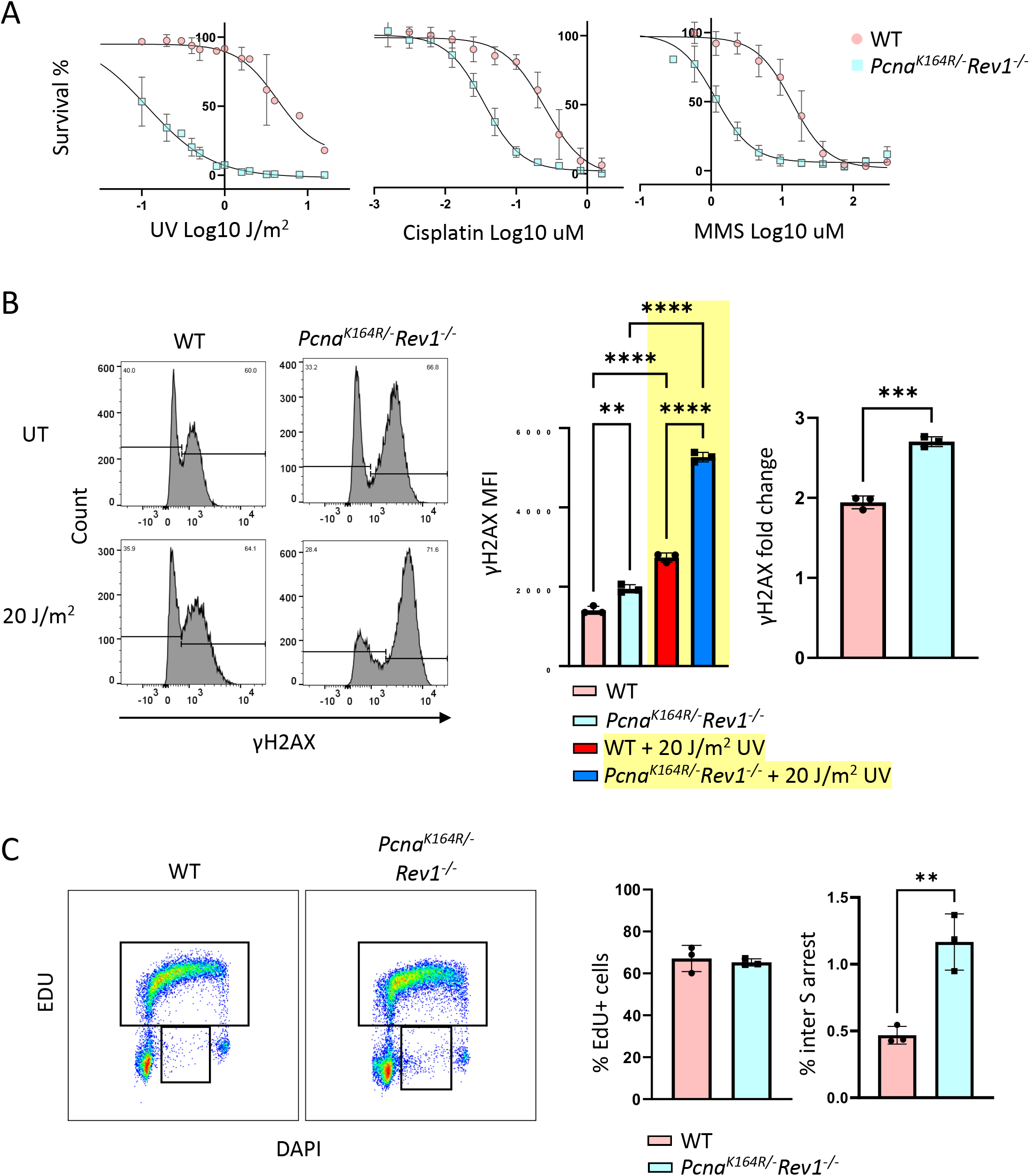
Consequences of DDT deficiency upon genotoxic stress and the cell cycle. (A) Survival curves of WT and DM lymphomas upon exposure of UV-C, cisplatin, or MMS. (B) Replication stress by mean fluorescent intensity (MFI) of γH2AX with or without UV-C exposure in lymphomas. (left) Histograms are from a representative experiment, (right) bar graph of the γH2AX MFI from the γH2AX positive population as indicated in the histograms. (C) Analysis and quantification of cell cycle progression via EdU incorporation in unperturbed WT and DM lymphomas.

To confirm that the hypersensitivity of the DM related to increased replication stress, we determined the levels of phosphorylated H2AX (γH2AX) as a marker for DNA damage and replication stress. Mean fluoresce intensity (MFI) of γH2AX was used as a relative measure of replication stress. Without treatment the levels of γH2AX in the DM were modestly increased compared to WT. In line with the hypersensitivity, treatment with a high dose of UV-C (20 J/m^2^) further increased replication stress 4 hours after exposure in both the DM and WT. Again, DM displayed much higher levels (Figure 1B, Supplementary Fig. 3A), indicating that the sensitivity to endogenous and exogenous genotoxins directly relates to the level of replication stress as measured by the accumulation of γH2AX.

As replication stress was increased in untreated conditions, we set out to determine the effect of a compound DDT defect on the cell cycle as determined by incorporation of the thymidine analogue EdU (5-ethynyl-2ʹ-deoxyuridine) in combination with the DNA stain DAPI. Somewhat surprisingly, a 20-minute pulse of EdU followed by flow cytometry failed to reveal a significant difference in the frequency of cells in S phase in DM (Figure 1C, Supplementary Fig. 3B). However, a significant increase of S phase cells that failed to incorporate EdU was found in the DM (Figure 1C). These cells likely activated an intra S checkpoint prior to the EdU pulse, resulting in an intra S phase arrest. Apparently, under endogenous stress conditions the cell cycle of most DM cells was unaffected, implying a similar rate of division but slight increase of intra-S aborted cells.

### Reliance of DDT deficient cells on CST

To investigate molecular dependencies and potential backup mechanisms of DDT in DM cells, we took advantage of WT and DM lymphoma cells to perform an unbiased whole genome dropout screen in unperturbed conditions (Figure 2A). Upon stable expression of Cas9 in DDT-proficient WT and DDT-deficient DM cells we used the YUSA V2 library (Tzelepis et al., 2016(55)), containing ≈79, 000 guides that target most murine protein-coding genes. We observed an excellent R^2^ of 0.7-0.8 between independent replicates in both WT and DM cell conditions, and moreover observed decreases of guides targeting essential genes over time, whereas non-essential genes remained unaltered (Supplementary Fig. 4A). Importantly, DM cells did not feature proliferation defects, which allowed us to directly compare both isogenic cell lines without confounding issues resulting from differential stress signals and replication defects. Amongst other interesting potential hits, two of the three components of the CST complex, TEN1 and CTC1 were readily identified, with the third component, STN1 (encoded by *Obfc1*) being just below the significance cutoff (Figure 2B). Interestingly, we did not find PrimPol, an important polymerase involved in repriming, suggesting that repriming capacity is not crucial for survival of DM cells.

**Figure 2.**
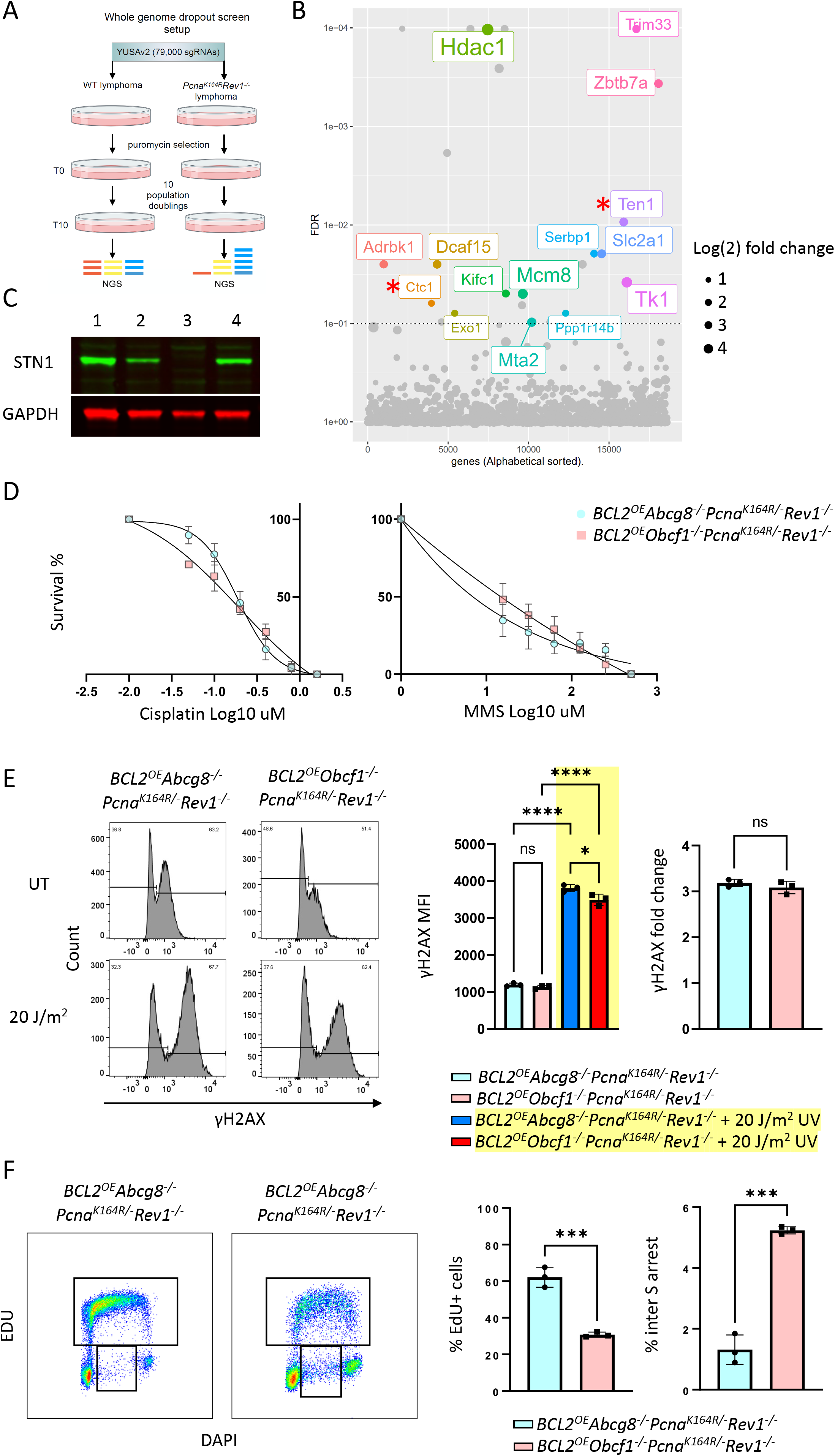
Investigation of the molecular dependency of DDT deficient cells. (A) Schematic overview of CRISPR-Cas9 whole genome dropout screen. (B) Overview of the results from the CRISPR-Cas9 whole genome dropout screen. (C) Western blot showing ablation of the STN1 protein in the lymphomas. 1) WT, 2) *Pcna^K164R/-^*;*Rev1^-/^*-, 3) *BCL2^OE^*;*Obcf1^-/-^*;*Pcna^K164R/-^*;*Rev1^-/-^*, 4) *BCL2^OE^*;*Abcg8^-/-^*;*Pcna^K164R/-^*;*Rev1^-/-^*. (D) Survival curves of STN1 KO and control lymphomas upon exposure of cisplatin or MMS. (E) Replication stress by MFI of γH2AX with or without UV-C exposure in STN1 KO and control lymphomas. (left) Histograms are from a representative experiment, (right) bar graph of the γH2AX MFI from the γH2AX positive population as indicated in the histograms. (F) Analysis and quantification of cell cycle progression via EdU incorporation in unperturbed STN1 KO and control lymphomas.

To validate the dependence of DM cells on CST, we opted for a STN1 knockout in the DM lymphomas, as STN1 specific antibodies were available. In line with the screen, we failed to derive a viable cell line. However, upon overexpression of the anti-apoptotic factor BCL2 in the DM setting we were able to establish a STN1 KO in these cells (Figure 2C). The BCL2 overexpression system allowed further exploration of the molecular relevance of CST in the DDT deficient cells. As a transduction control the *Abcg8* gene was used, a gene commonly used as control gene in whole genome CRISPR screens. When exposed to cisplatin or MMS, CST deficient DM cells did not show a marked increase in sensitivity compared to the BCL2 overexpressed control (Figure 2D, Supplementary Fig. 4B, 4C). This suggests that CST is not directly involved in DDT of these specific exogenous lesions.

In support of this notion, the MFIs of γH2AX were found comparable between the BCL2 control and its STN1 KO (Figure 2E). A slight decrease in replication stress was found upon UV treatment, though comparison of the fold-changes seems to negate this difference. The level of replication stress correlated with drug sensitivities (Figure 2D, 2E).

Strikingly, using EdU to investigate the cell cycle profile revealed drastic changes in the STN1 KO (Figure 2F). The number of EdU positive cells was severely reduced, the number of EdU negative inter S phase arrested cells was found increased (Figure 2F). This suggests a role of CST in stabilizing replication during endogenous stress.

Taken together, unlike DM cells, the changes in the cell cycle of the STN1 KO are not reflected by an increase in replication stress. The primary role of CST in prohibiting replication stress appears to relate to endogenous replication impediments rather than coping with specific exogenous lesions.

### Efficient repriming increases fork speed in DDT deficient cells

As DDT is primarily demanded and activated during genome replication and DM displayed increased replication stress, we studied replication fork progression using the DNA fiber assays to visualize and analyze individual replicons (Figure 3A)(56). Under standard culture conditions, we observed a significant increase in the replication speed in DM lymphomas (Figure 3B). This likely relates to repriming upon encountering a lesion, as one of the few options left to the DM cells. Upon UV-C exposure, a decrease in fork speed in the DM was observed, while an increase was found in WT cells. This decrease in the DM setting implies an increase in fork stalling caused by the inability to effectively tolerate the high abundance of UV-C induced DNA lesions. In the WT setting, the increase in fork speed implies an increase in repriming.

**Figure 3.**
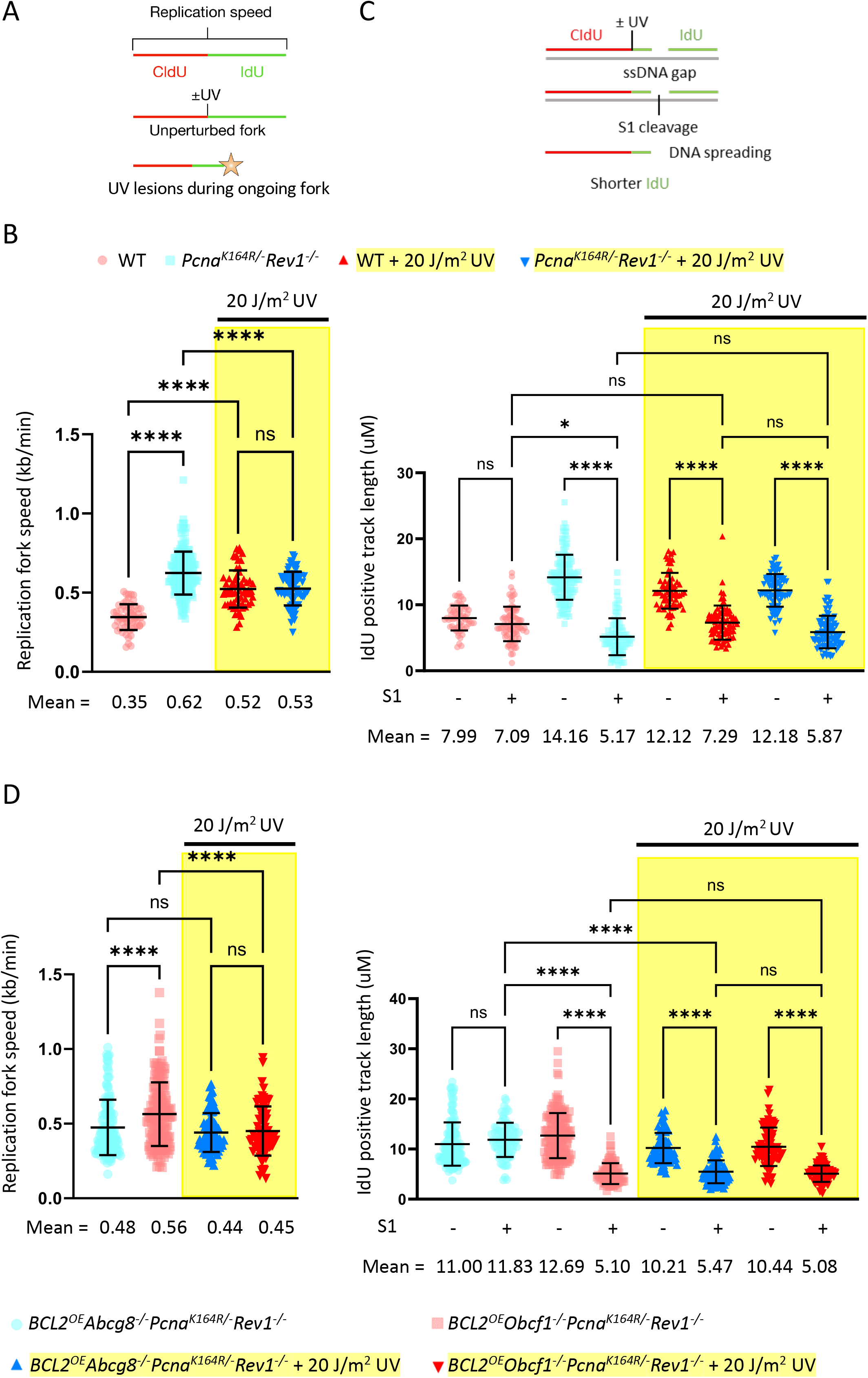
Replication fork speed and S1 nuclease assay in DDT deficient lymphomas and STN1 KO. (A) Schematic representation of the DNA fiber labeling experiment. The upper panel represents the two labeling periods and ongoing replication fork without encountering exogenous lesions. The lower panel illustrates the replication fork encountering a UV-C induced lesion. (B) (Left) Replication speed, lymphomas were first labelled with CldU for 20 minutes and then exposed to 20 J/m^2^ UV-C or mock treatment, followed by 60 minutes labelling with IdU. The lengths of IdU tracks were measured and the average replication speed (kb/min, error bar indicates standard deviation) is displayed. (Right) IdU track length with or without S1 treatment and UV-C. Same labeling and treatment strategy was used as for replication speed assays. *p* values are based on Kruskal-Wallis test, * P<0.05, ** P<0.01, *** P<0.001, **** P <0.0001. (C) Schematic representation of the DNA fiber labeling experiment with S1 nuclease digestion. In the upper panel a ssDNA gap is illustrated in the second label (IdU). After S1 nuclease treatment, in the lower panel, the ssDNA gap is cleaved, and a shorter IdU track length is left. (D) Same as in B but with the STN1 KO lymphoma and control.

If a system becomes reliant on repriming, single strand gaps are expected to be generated. In DNA fibers, these ssDNA gaps can only be detected and quantified indirectly by an S1 nuclease treatment prior to spreading (Figure 3C). Applying this approach, we observed a profound decrease in track length in the DM lymphoma, while WT cells remained unaffected under standard culture conditions (Figure 3B). These findings indicate that the observed increase in fork speed likely relates to increased repriming. Upon UV-C exposure and S1 treatment only WT cells displayed decreased track length as compared to non-UV-C exposed but S1 treated conditions, while the track length in DM cells remained the same (Figure 3B). This implies that the increased replication speed in WT is based on increased repriming activity, which becomes necessary when challenged with exogenous replication stressors. In contrast, the DM system remains reliant on repriming in both, unchallenged and challenged conditions (Figure 3B), suggesting that DM predominantly use repriming as a mode DDT to alleviate replication stress(57).

Our results indicate that a CST deficiency in the DM did not result in a major defect in handling exogenous genotoxic stress but exerts its function primarily under specific endogenous stress conditions, as shown by cell cycle data (Figure 2F). To study this more directly, we determined the consequences of a ST1 KO on replication fork speed under endogenous and exogenous replication stress conditions (Figure 3D). Here, we took advantage of BCL2 overexpression to study the CST deficiency in the absence of a global DDT. Unexpectedly, BCL2 overexpression appeared to reduce the increased replication speed in the DM. In the absence of STN1 fork speed was found increased. Upon induction of UV lesions, the lack of a CST complex had no additional effects on top of what was observed in the DM (Figure 3B, 3D). Using the S1 cleavage assay we determined the extent of ssDNA in nascent DNA fibers in these cells (Figure 3D). Remarkably, while BCL2 overexpression in the DM reduced the amount of ssDNA, the STN1 KO displayed a marked decrease in track length. This indicates a critical contribution of the CST complex in promoting fork stability and preventing the generation of ssDNA during replication under endogenous conditions. Again, upon UV-C exposure there was no additional changes of a STN1 KO DM as compared to the phenotype of the BCL2 overexpressed DM control, suggesting that CST activity prefers specific types of lesions or impediments(58, 59).

### DDT-deficient cells are prone to acquire large genomic deletions

To investigate the role of DDT in mutagenesis and genome maintenance we performed WGS in DM lymphomas under endogenous and exogenous stress conditions. Analyzing the large-scale mutations (structural variances), we observed a significant increase in deletions larger then 50bp in the DM (Figure 4A). No significant changes were detected in the other types of large-scale mutations such as large insertions or duplications, which in general are molecularly more complex and hence are less frequent mutational events. To examine the deletion landscape in DDT-deficient cells in more detail, we plotted the size distribution of deletions and identified three distinct peaks of progressively larger deletion size (termed type 1, 2, and 3 deletions) (Figure 4B). Importantly, type 3 deletions, ranging in size from 0.4 to 4.0 kbp, were found to be increased in DM cells compared to WT cells (Figure 4B). To determine if the formation of these deletions could also be induced through exogenous lesions we exposed the cells to UV-C irradiation. Compared to WT, the DM cells showed a specific increase in type 3 deletions upon UV-C (Figure 4C). Notably, when UV exposed DMs were compared against non-treated DMs we also saw a selective increase in these deletions, showing that these large deletions are damage-inducible in DDT deficient cells (Figure 4D). Overlaying the total number of deletions in all conditions demonstrates a role of DDT in preventing type 3 deletions upon encountering exogenous lesions (Figure 4E). No difference was observed in the genomic location of type 3 deletions seen between WT and DM cells untreated or treated with UV (Supplementary Fig. 5A, 5B). This argues against specific sites of genome that are more prone for type 3 deletions.

**Figure 4.**
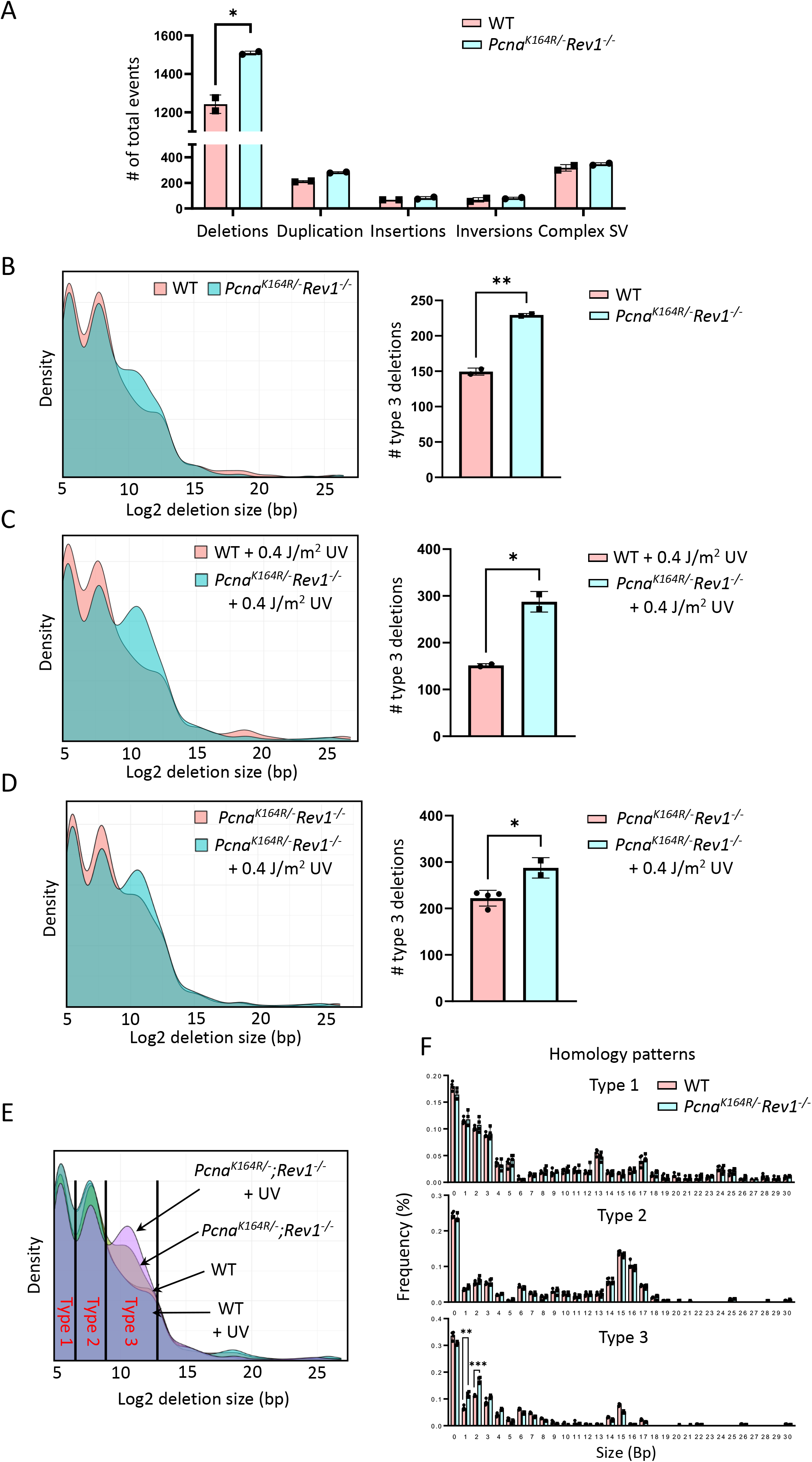
Whole genome sequencing analysis of DDT deficient cells. (A) Number of structural variances found in WT and DM lymphomas. *p* values were determined using an unpaired student t-test. * P < 0.05. (B) (Left) Deletion size density plot of WT and DM lymphomas, average of two samples. (Right) Number of type 3 deletions found in WT and DM lymphomas. *p* values were determined using an unpaired student t-test. ** P < 0.005. (C) (Left) Deletion size density plot of deletions that occurred after the exposure of 0.4 J/m^2^ UV-C of the WT and DM lymphomas, average of two samples. (Right) Number of type 3 deletions found in WT and DM lymphomas after UV-C exposure. *p* values were determined using an unpaired student t-test. * P < 0.05. (D) (Left) Deletion size density plot of deletions that occurred in untreated DM lymphomas compared to DM lymphomas after the exposure of 0.4 J/m^2^ UV-C. Average of four samples in untreated conditions, average of two samples in UV-C treated conditions. (Right) Number of type 3 deletions found in WT and DM lymphomas under indicated conditions. *p* values were determined using an unpaired student t-test. * P < 0.05. (E) Deletion size density plot of deletions all deletions present in the cultured WT and DM cells and the UV exposed WT and DM cells with the different type of deletion ranges indicated. (F) Bar graph of the percentage of homology found at indicated types of deletion ranges of WT and DM lymphomas. *p* values were determined using multiple unpaired student t-tests with Two-stage step-up correction (Benjamini, Krieger, and Yekutieli). ** q < 0.005, *** q < 0.0005.

Given the recurrent, characteristic, multimodal size distribution of genomic deletions and that they likely require at least a single DSB(60), we hypothesized that each modus underlies a molecularly distinct DSB processing pathway. To approach this hypothesis, we examined the extent of microhomology at the junction sites of the deletions. Consistent with this concept, each peak displayed unique microhomology preferences at the break/junction sites (Figure 4F, Supplementary Fig. 5C), suggesting that different molecular processes are involved in generating this characteristic size spectrum of genomic deletions. Type 1 deletions, ranging from 30-90 bp, displayed varied pattern of homology sizes with an increase in microhomology of 1-3 bps. Interestingly, the type 2 deletions ranging between 90-500 bp showed a very specific 14-17 bp micro-homology preferences. Type 3 deletions, with the specific increase in the DDT-deficient cells, shows again an increase in microhomology of one and two base pairs under endogenous conditions. Unique about the homology pattern in the third peak, was that the frequency of 1-2 bp microhomology was increased in the DM cells compared to the WT cells, whereas the homology pattern of all other base pairs was found non-significantly changed between genotypes (Figure 4F). UV exposure only affected the homology pattern of type 3 deletions, where the DM cells had a significant increase in deletions with a microhomology signature of 3 bp compared to the non-treated DM cells (Supplementary Fig. 5A). These results support the concept that the multimodal size-distribution of genomic deletions underlies distinct molecular activities where DNA repair associated with type 3 deletions likely relates to one of the yet poorly defined A-EJ pathways. This argues against the involvement of canonical NHEJ, single strand annealing (SSA), and HR generating type 3 deletions(61, 62). Taken together, our results reveal that DDT prevents the accumulation of large genomic deletions in a DNA damage-dependent manner. These large deletions are likely repaired via a distinct, yet undefined, microhomology dependent A-EJ pathway(11).

Surprisingly WGS results of *Pcna^K164R/-^;Rev1^-/-^* cells failed to reveal small-scale mutations (Supplementary Fig. 6). These data indicated that the frequency and distribution of point mutations were highly comparable. Likewise, the frequency and distribution of InDels, and the mutational spectra were highly similar between the WT and DM clones (Supplementary Fig. 6B-6E). As only the removal of DDT did not reveal any major differences in these types of small mutations, and as DDT is responsible for resolving different types of lesions, we examined how UV-C affected genomic stability of DDT-deficient cells. In line with the previous results, no differences were observed in substitutions and InDels (Supplementary Fig. 6). Furthermore, the single base pair substitution spectrum as described by Alexandrov *et al*.(63) failed to reveal significant changes in the type or quantity of point mutations. Despite the critical role of DDT in tolerating DNA lesions and other structural impediments during replication, our results argue that replicative bypass or persistent replication blockade in the absence of DDT, does not result in major changes in small scale mutations. Apparently, DDT plays a major role in alleviating replication-stress induced genome instability by preventing type 3 deletions, without introducing small scale mutations.

### Type 3 deletions can dominate the deletion landscape in human tumors

If conserved evolutionarily, it is expected to find a similar multimodal size distribution of genomic deletions in WGS samples of different species. To address this question and determine whether type 3 deletions arise in human cells as well, we took advantage of the publicly available WGS database from the Hartwig Medical Foundation (HMF). This unique database contains more than 4, 500 human tumors. Random sampling of tumors and averaging the deletion size distributions revealed -except for type 2 deletions-a similar size distribution of human tumor samples and the mouse lymphoma (Figure 5A). Deletions within the smallest range of size distributions (30-75 bp, type 1) occurred in both mouse and human tumors whereas the type 2 deletions (120-360 bp) were only detected in the mouse lymphoma. Interestingly, a recurrent proportion of type 3 deletions (0.4-4.0 kbp) was present in most human tumor types (Figure 5A). This recurrent multimodality suggests evolutionarily conserved molecular mechanisms underlying the size-restrictions of these deletions. Furthermore, when compared to randomly selected samples of the same tumor type a subset of tumors was found highly enriched in type 3 deletions, indicating the existence of conditions favoring type 3 deletions in human tumors (Figure 5A). To determine the frequency of human tumors containing type 3 deletions we looked at the percentage of deletions that fall within the range of type 3 deletions compared to the total amount of deletions present. Consistently across different tumor types, 20 to 35 percent of tumors seem to contain 25 percent or more type 3 deletions (Figure 5B). Taking this 25-percent cutoff and plotting the density profiles, clearly indicated a sizeable increase of type 3 deletions independent of tumor origin (Figure 5C).

**Figure 5.**
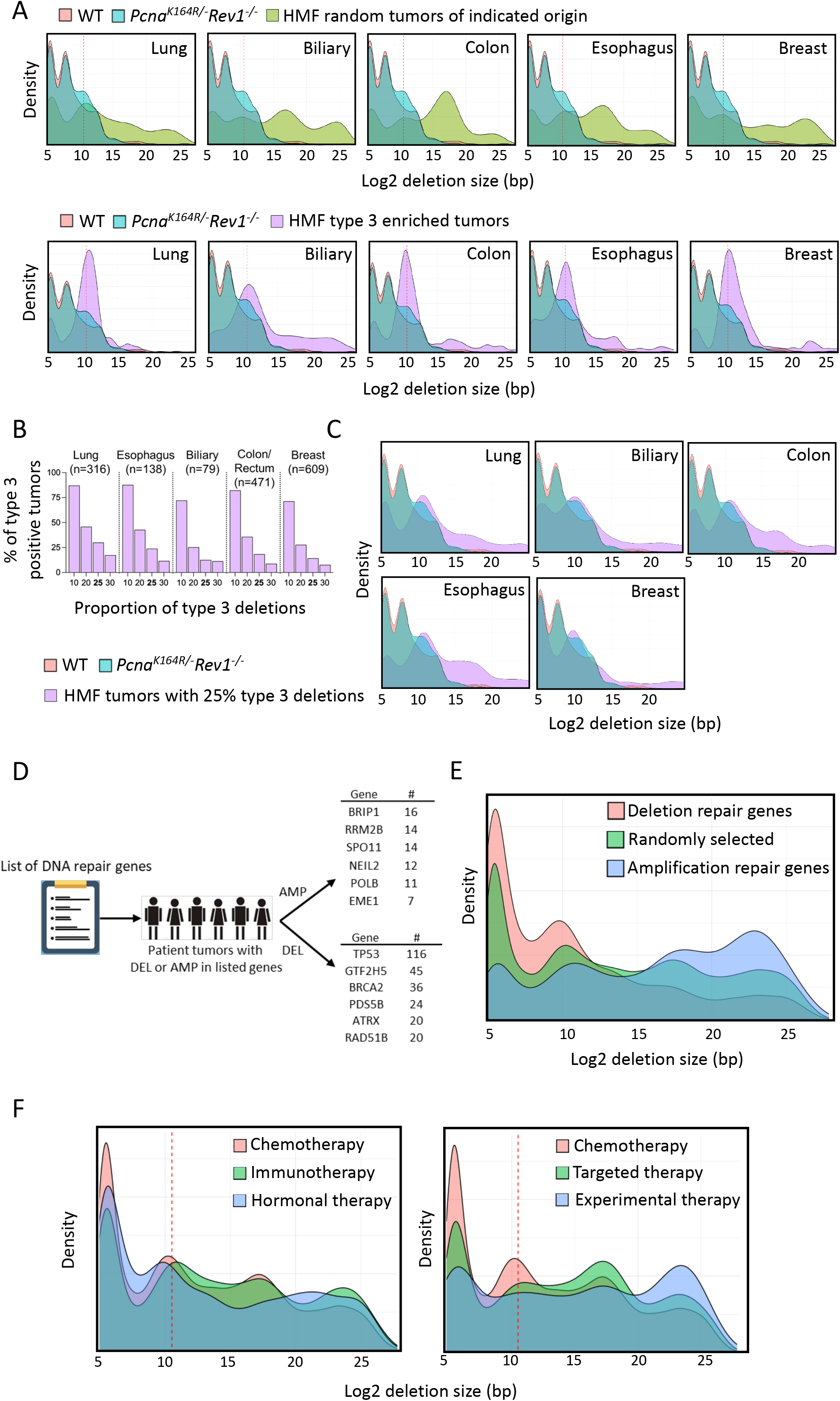
Analysis of large deletions in human tumors. (A) (Top) Combined deletion size density plot of 50 randomly selected human tumors of the indicated tumors type including WT and DM mouse lymphoma. (Bottom) Combined deletion size density plot of a selection of human tumors that are enriched in type 3 deletions of the indicated tumor type including WT and DM mouse lymphoma. (B) Frequency of tumors with the indicated percentage of all deletions being within 0.4-4 kbps. (C) Deletions size density plot of tumors with 25% of their deletions being within 0.4-4 kbps. (D) Scheme to investigate the effect of DNA repair gene proficiency on deletion sizes. Patients tumors were selected with either a deletion or 20-fold amplification of listed DNA repair genes. The top 6 genes with the most tumors found with the indicated mutation are shown and used for the analysis. (E) Combined deletion size density plot of tumors with either an amplification or deletions in DNA repair genes or randomly selected tumors. (F) Deletion size density plots of tumors, regardless of tumor type, that received different types of treatments.

### DNA repair defects and activities correlate with the type 3 deletions in human tumors

There is a delicate balance between DNA repair activity and defects in determining whether mutations such as deletions are acquired. As replication stress can provoke DSBs which underly the formation of genomic deletions, we investigated the role of different DNA repair genes in the generation of type 3 deletions in human tumors. Tumors were stratified based on either a deletion or 20-fold amplification of known DNA repair genes. Tumors with the six most frequently amplified or deleted genes were compared (Figure 5D). The combined density profiles of the deletion sizes of these tumors revealed a remarkable trend where the loss of DNA repair genes correlated with an increase in type 3 deletions whereas an amplification of DNA repair genes led to their decrease (Figure 5E, Supplementary Fig. 7A, 7B). As no specific relation was found between these genes and specific DNA repair pathways, distinct DDR defects are likely to enhance replication stress and DSBs, kick-starting the generation of type 3 deletions as well as others. The fact that except for type 2 deletions the deletion profiles between human tumors and mouse lymphomas appear similar, along with the reliance of these types of deletions on the proficiency of DNA repair genes in both species, implicates a conserved molecular mechanism and/or (epi)-genetic condition underlying the generation of type 3 deletions. Furthermore, comparing different therapies showed that the deletion spectra and specifically the frequency of type 3 deletions are influenced by the treatment modality (Figure 5F). A preference in the accumulation of type 3 deletions was found upon chemotherapy, immunotherapy, and hormonal therapy, while decreased in experimental and targeted therapy especially in tumors treated with Tamoxifen, Durvalumab, Imatinib, or Lorlatinib (Figure 5F, Supplementary Fig. 8)

## DISCUSSION

Both, PCNA-Ub and REV1 are critical components of the DDT system. Compound mutants lacking PCNA-Ub and REV1 were found synthetic and embryonic lethal (PNAS, Supplementary Fig. 1A). This synthetic lethality could be rescued by P53 downmodulation or inactivation. Taking advantage of this knowledge, we here investigated the molecular dependencies and genomic impact of a global DDT defect in the mammalian system. As expected, DDT deficient DM cells were hypersensitive to replication blocking lesions and in accordance displayed a marked increase in replication stress responses. Despite these impairments, the cell cycle remained largely unaffected under unperturbed conditions, but with a noticeable increase of cells arrested in inter S phase. The fact that cells can survive in the absence of PCNA-Ub and REV1 dependent DDT suggest the existence of a compensatory DDT backup mode. To identify molecular dependencies of DM cells, we performed a whole genome CRISPR-Cas9 drop out screen. Interestingly, the CST complex was identified as essential for DM cells. Moreover, as CST has been associated with replication, including fork stability, telomere maintenance, repriming via POLA, tolerance of oxidative damage, as well as de novo origin firing, we characterized the contribution of CST in alleviating replication stress under compromised DDT conditions(39, 40, 45, 58, 59). Further validation by a KO of STN1 (encoded by *Obfc1*) confirmed a vital dependence of DM cells on CST. To enable the investigation of a ‘dropped out’ CST target complex, we first had to overexpress the antiapoptotic BCL2 by lentiviral transduction ensuring cell survival. In the absence of CST, DM lymphoma cells were found similarly sensitive to exogenous stressors as CST proficient DM controls, whereas under non-challenged conditions the cell cycle was severely affected. This argues against an important function of CST response to exogenous stressors. Instead, CST appears primarily enrolled in processing endogenous lesions and other replication impediments such as oxidative damage, G4-stacks, hairpins, and Z-DNA structures(58).

As DDT is essential for mammalian replication and cell proliferation which can be rescued by p53 inactivation, we studied the effect of a DM on replication using the DNA fiber assay(56, 64, 65). A global DDT defect resulted in a highly increased replication fork speed, which is consistent with a report on the impact of replication stress on fork speed and repriming(66, 67). Indeed, as suggested by S1 nuclease digestion of nascent DNA fibers, repriming appears to predominate as a backup DDT mode to bypass lesions in the absence of PCNA-Ub and REV1 and prohibit cell death by prolonged fork stalling(57). Furthermore, UV-C exposure decreased the fork speed in the DM while increasing it in the WT. The increased fork speed in WT cells likely relates to repriming as revealed by the shortening of the track length upon S1 treatment. In line, the decreased fork speed in the DM is probably caused by increased fork stalling due to an impaired capacity to process the excessive number of lesions leading to an intra S phase arrest and cell death (Figure 1C). The S1 data indicate that also under UV-C conditions DM cells rely on repriming to warrant fork progression.

Remarkably, as shown in (Figure 3D) overexpression of BCL2 in DM cells modulated DDR signaling such that besides a survival advantage, sufficient replication fidelity is gained to reduce repriming (Figure 3D, right panel, light blue) and associated fork speed (Figure 3D, left panel, light blue).

In support of previous conclusion, STN1 KO had no effect other than those observed in the DM cells, when exposed to UV. This suggests that CST is primarily required in alleviating endogenous replication impediments. Interestingly, our STN1 KO data revealed an increased fork speed (Figure 3D, right panel, light red) accompanied by increased repriming activity (Figure 3D, right panel, light red), implicating that CST exhibits its contribution to DDT activity by stabilizing replication forks and prohibiting repriming. Failure to prohibit repriming in DM cells lacking CST likely explains the dependence of DM on CST i.e., DM cells must prevent extensive repriming to warrant survival. This interpretation is in line with the observation, that PrimPol, a potent primase at stalled forks was not found in our drop out screen of DM lymphoma cell line(57, 68). In this way fork stabilization may facilitate an inefficient POLD/E polymerase switch to TLS polymerase by unmodified PCNA, as suggested previously(69, 70).

A central remaining question in the DDT field relates to the pro- and anti-mutagenic contribution of DDT pathways to genome maintenance. Applying WGS we here identified a critical role of DDT in preventing the generation of damage-inducible large genomic deletions spanning between 0.4-4.0kbp, which were also found in human tumors, which in general experience replication stress(71, 72). Furthermore, breakpoint junction analysis revealed that the generation of these deletions prefers a 1 to 3 base pair microhomology. Notably, the size density profile of deletions in the mouse cells revealed a multimodal distribution defined as type 1, 2 and 3 deletions, characterized by specific size and microhomology preferences. The first type of deletions is reminiscent of the small deletions previously identified by TMEJ(73) and depend on single bp microhomologies. In contrast to these type 1 deletions which occur in mice and humans, type 2 deletions were specific for the murine system. Type 2 deletions are characterized by unusual microhomology preference of 15 and 16 bp. Taken together, the discrete size preferences associated with distinct patterns of microhomology at the break-junction sites, supports a concept where distinct molecular activities underlie the generation of these deletions.

Interestingly, an unbiased analyses of whole genome sequences of human tumors revealed the presence of similar sized type 3 deletions. Type 3 deletions were highly abundant in all tumor types. Replication stress is a hallmark of tumors(71, 72) and in line with this notion these type 3 deletions correlated with DNA repair status. Amplification of repair genes prevented type 3 deletions and inactivation of DNA repair genes favored their generation. Furthermore, the treatment modality of tumors was found to affect the deletion size spectrum independent of the tumor type. Based on these results, we favor a model where prolonged fork stalling leads to fork collapses and associated DSBs. Besides replication stress, deletion initiated by DSBs may also arise through other alterations in the DNA damage response network, suggesting DSBs as critical intermediate towards type 3 deletions. Alternatively, deletions may arise by looping of nonreplicated tracks of ssDNA independent of a DSB, as suggested recently(74). Long tracks of single stranded underreplicated DNA, spanning up to several kilo base pairs were reported by *Lopes et al*. in yeast(75), correlating with the size of type 3 deletions, hinting that under-replication may underlie these deletions.

A recent report indicated that deficiencies of single TLS polymerases in *C. elegans* resulted in the formation of large 100-600 bp genomic deletions in a POLQ-dependent manner(76). Though these deletions display both, a different size and microhomology pattern, it argues in favor of DDT preventing large deletions in multiple phyla of life. Likewise, point mutagenesis in TLS-deficient animals was also not observed(76), in line with the absence of overt point mutagenesis in our lymphoma lines. Additionally, large deletions (>50 bp) were detected in HSCs and p53-immortalized cells taken from *Fancd2^-/-^Aldh2^-/-^* mice(2). These findings are consistent with the concept where unresolved lesions, causing an increase in replication stress, can induce large deletions. The link between ICL repair and large deletions can give an explanation of the existence of type 3 deletions in human tumors, as these tumors frequently lose ICL repair and are often treated using DNA damaging agents(77). Consequently, ICLs will accumulate leading to replication stress and DSBs that serve as possible substrates of type 3 deletions. This concept is supported by the result that chemotherapy favors type 3 deletions. Interestingly, the mutagenic signatures of primary and p53-immortalized *Fancd2^-/-^ Aldh2^-/-^* cells appear very similar(2), arguing against a general role of P53 on the mutagenic landscape and in extension in our experimental settings. Additionally, similar findings such as the absence of small scale mutations and formation of large deletions in *C. elegans*(78), a non-transformed system, also oppose a direct role of P53 in the formation of these large deletions.

To conclude, this study firmly establishes a critical non-redundant contribution of PCNA-Ub and REV1 in DDT. An unbiased genome wide CRISPR screen identified the CST complex as essential back up in the absence of PCNA-Ub and REV1 facilitated DDT, where CST appears essential in preventing excessive repriming. Furthermore, we demonstrate that DDT exerts an anti-mutagenic function within the DDR network, by alleviating replication stress to protect the genome from structural variances. A multimodal size distribution of genomic deletions was observed. This multimodality is characterized by distinct MH signatures at the break ends. Our results indicate that DDT plays a critical role in preventing large deletions of the third modality, here classified as type 3 deletions. The multimodality likely reflects distinct stress conditions and repair activities, where type 3 deletions are a signature of replication stress. Future work will have to delineate the molecular conditions and determinants underlying the deletion mechanisms.

Remark: While finalizing this report, Z. Gyüre et al. (79) reported very similar interesting findings on the role of DDT in preventing large genomic deletions. The fact that both data sets were gathered and interpreted independently, highlights the relevance of both studies.

## Supporting information

All Suppl. Figures

## ACKNOWLEDGEMENTS

The authors wish to acknowledge R. Bin Ali from the Mouse Clinic for Cancer and Aging research (MCCA) Transgenic Facility for his help in generating Rev1^-/-^ mouse, Roel of Pruntel for sequencing, This work was supported by grants from the Dutch Cancer Society (KWF) to H. Jacobs (KWF NKI-2016-10032, and KWF NKI-2017-10796).

## CONTRIBUTIONS

DdG, AS, RS, CL, RLB and HJ designed the study and interpreted the results. DdG, AS, RS, MK, BM, PCMvdB, OAB, BP, SO, JJIC, MA, CL performed experiments. DdG, AS and HJ wrote the manuscript. HJ wrote the projects.

## COMPETING INTEREST

The authors do not have any competing interest to declare.

---

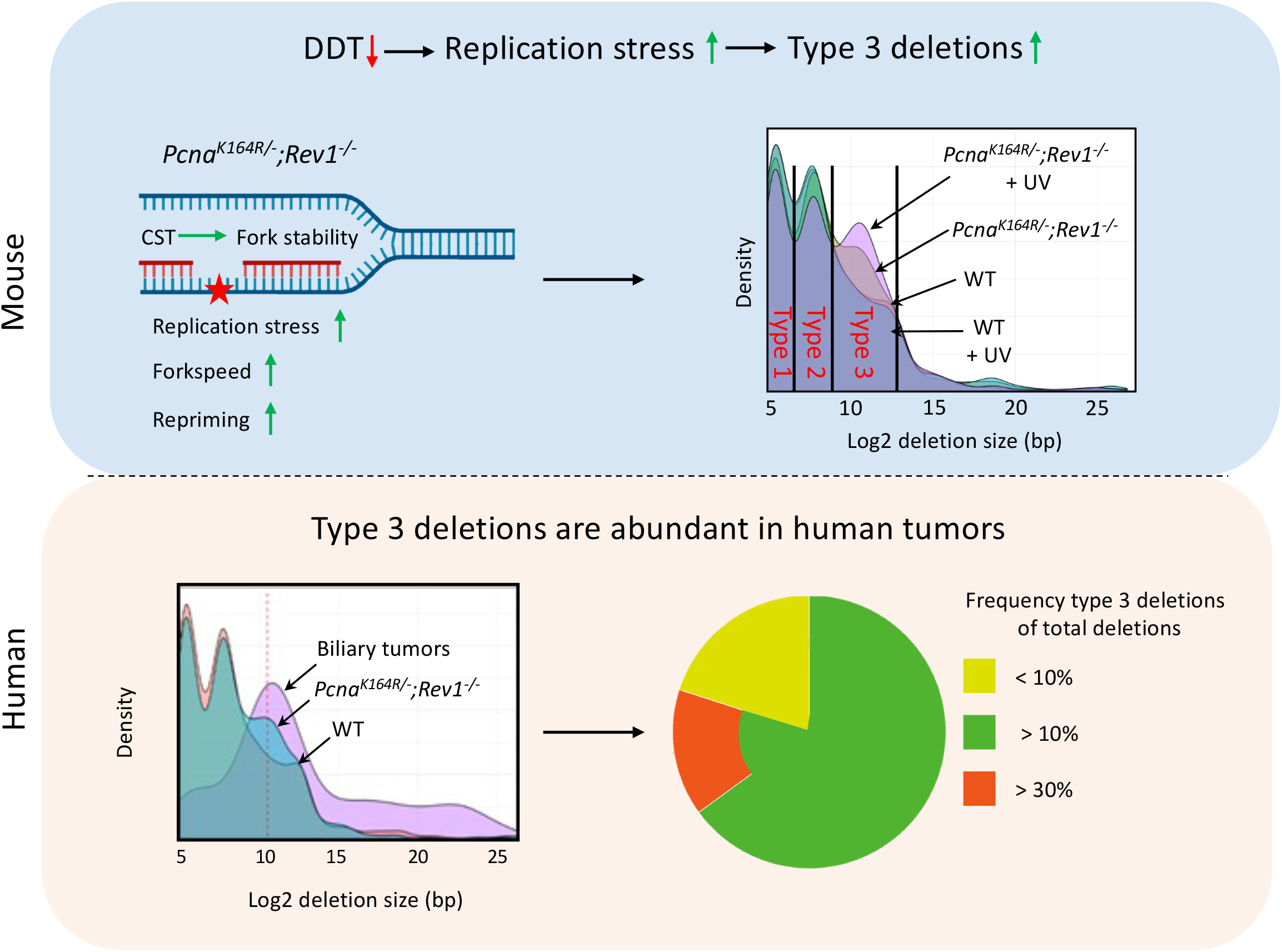

## Notes

### Competing Interest Statement

The authors have declared no competing interest.

### Summary of Updates

Corrected some typographical errors and improved layout.

