## Supplementary material for "Molecular dependencies and genomic consequences of a global DNA damage tolerance defect": All Suppl. Figures

Supplementary figure 1

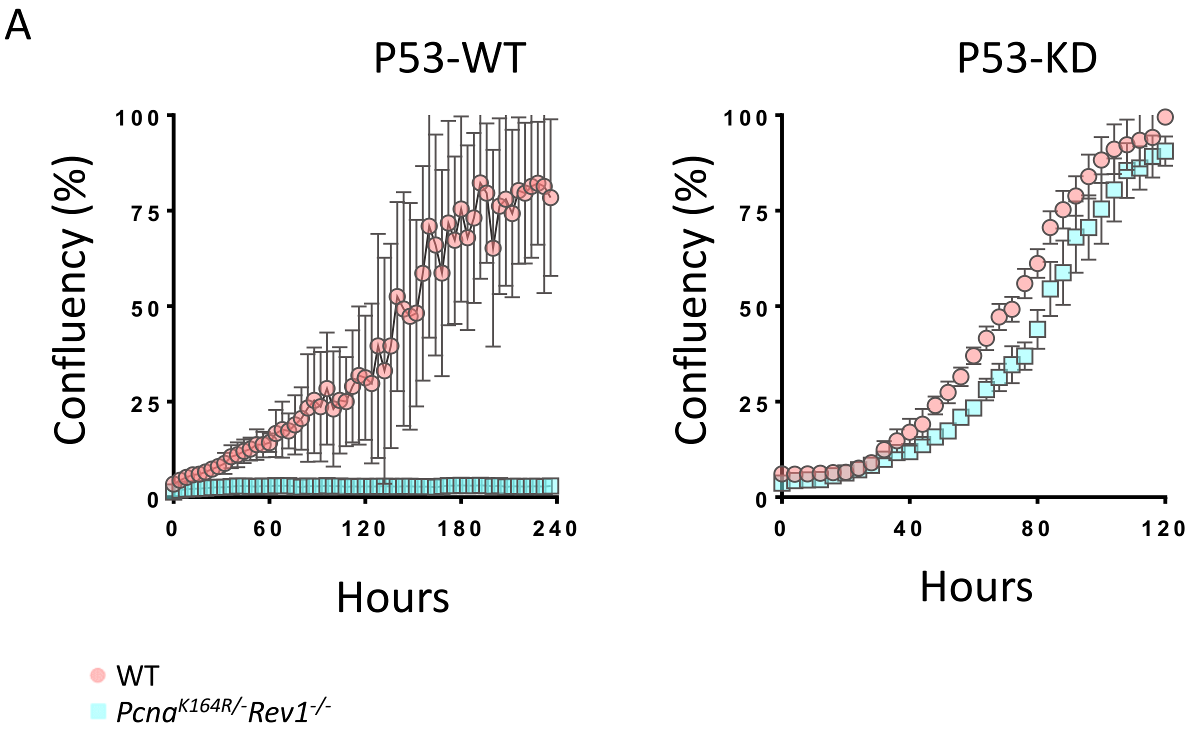

### Supplementary figure 2

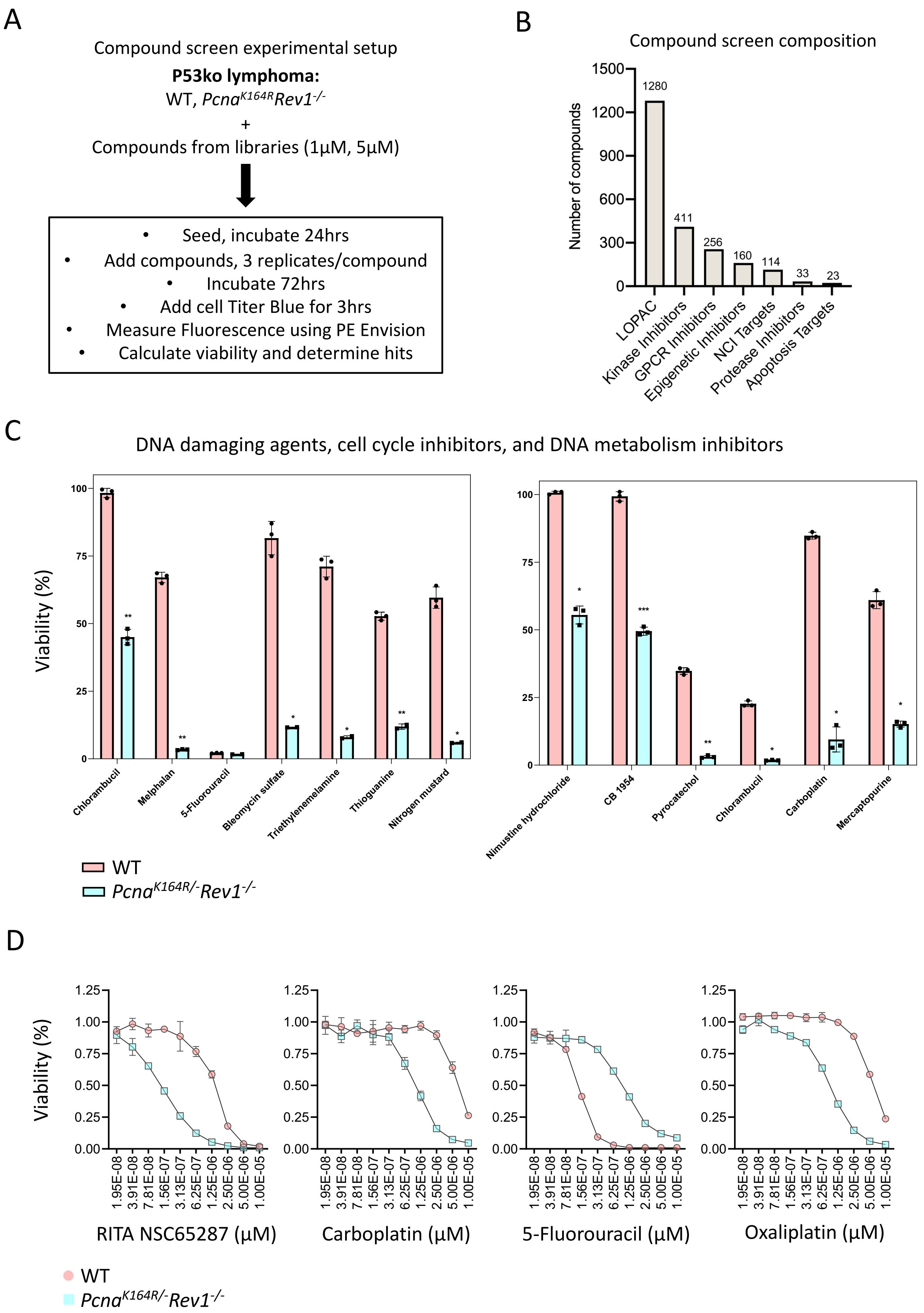

### Supplementary figure 3

A

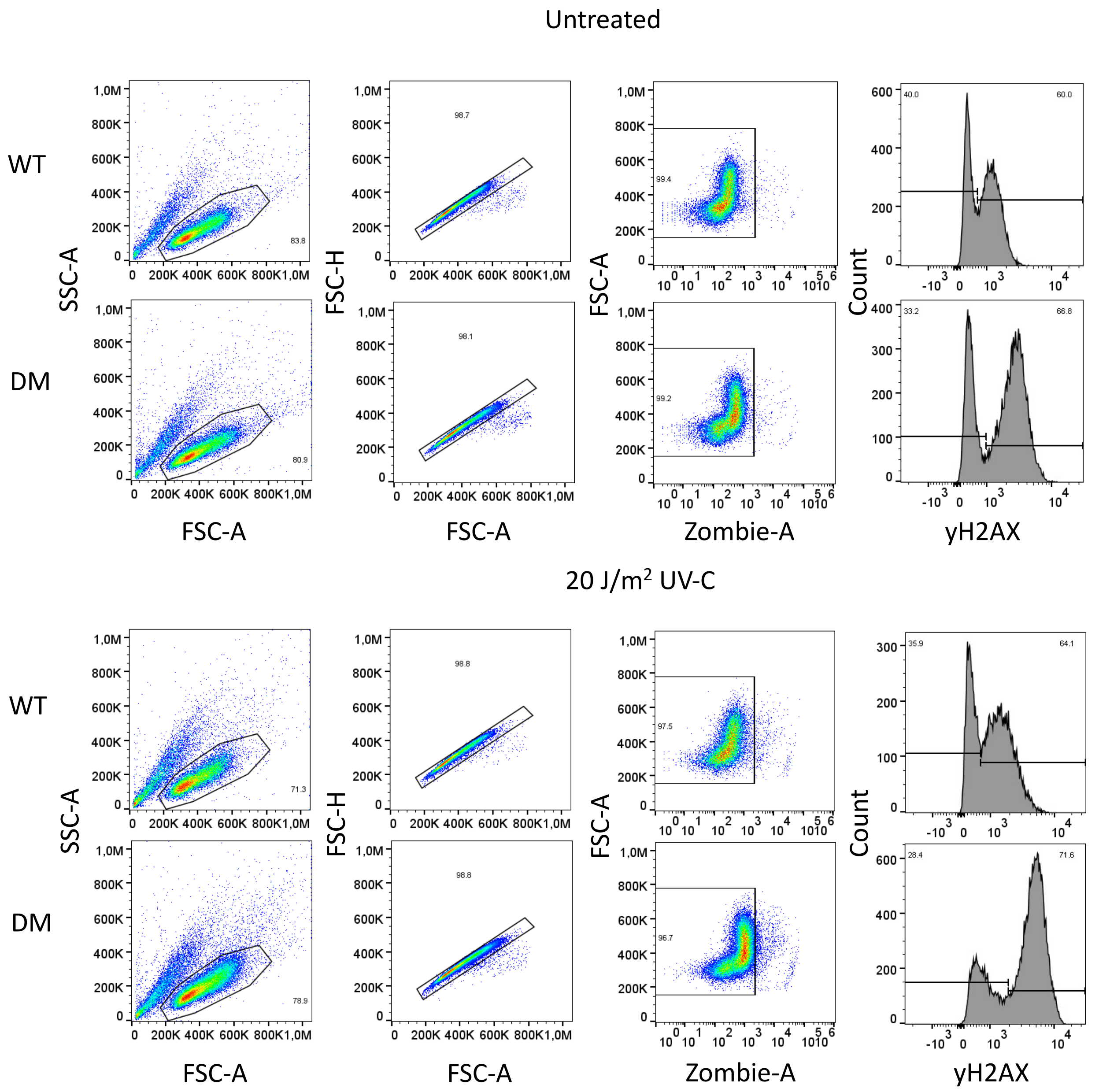


B

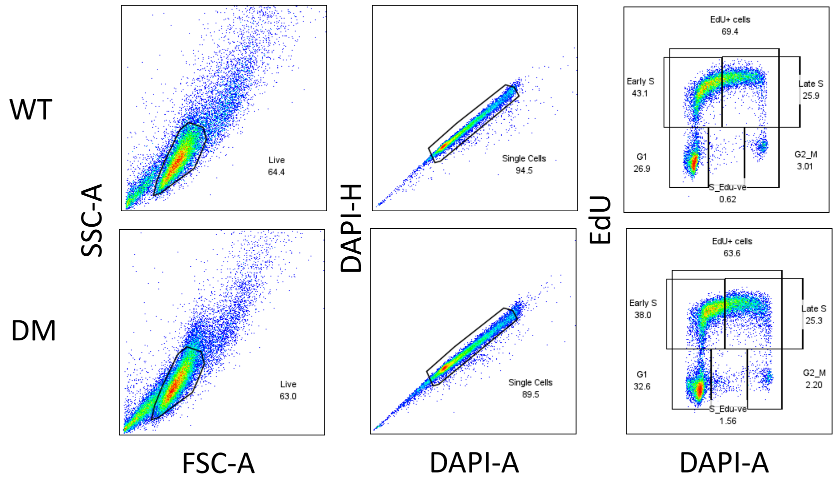

### Supplementary figure 4

A

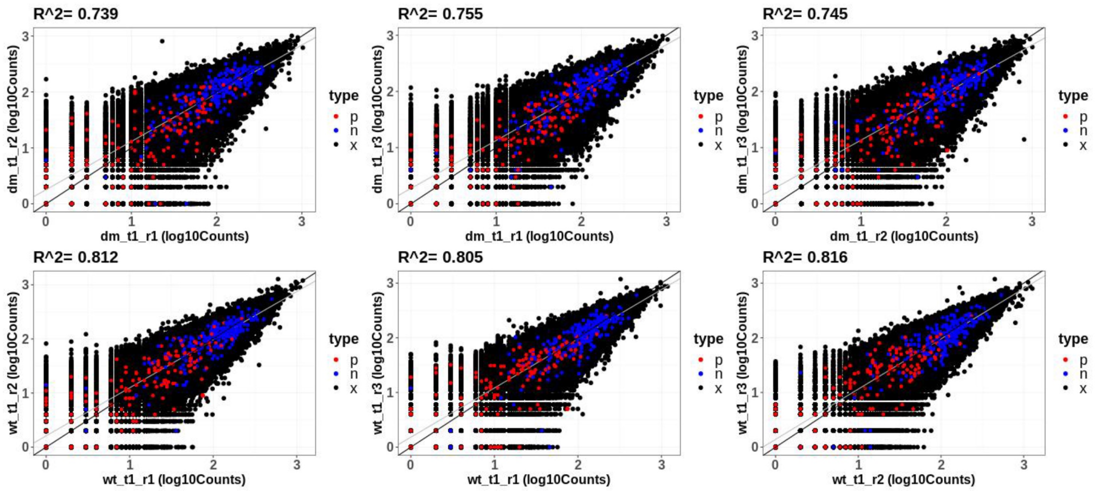


B

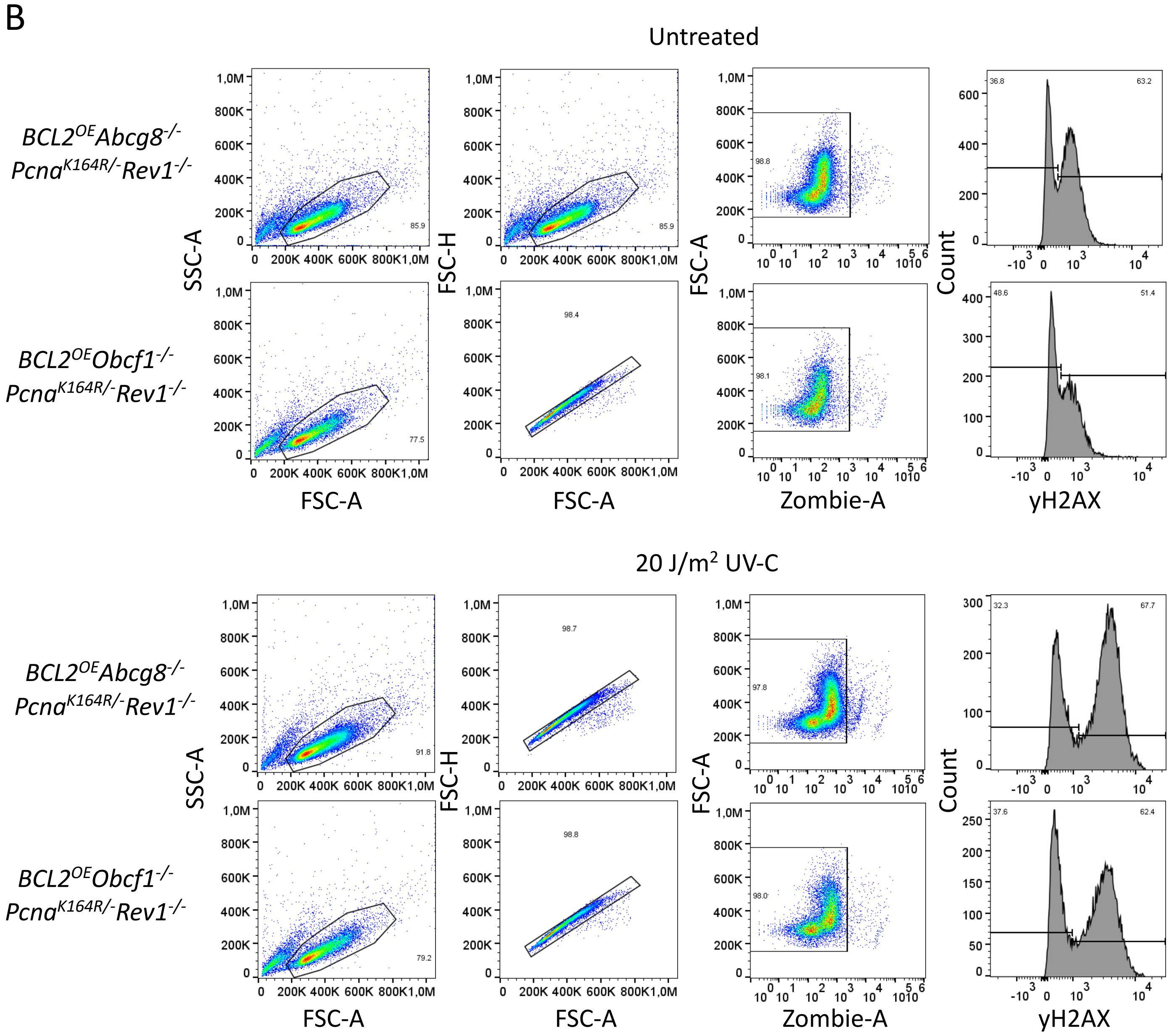


C

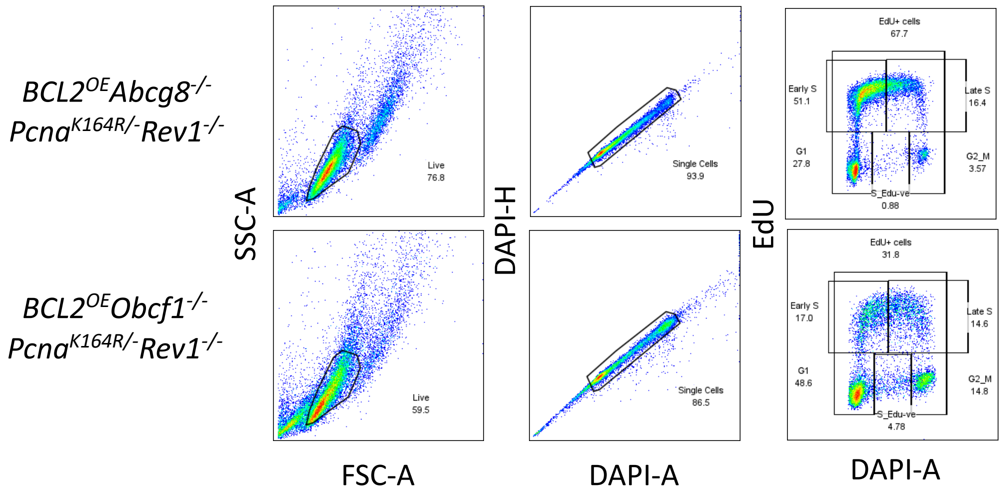

### Supplementary figure 5

A

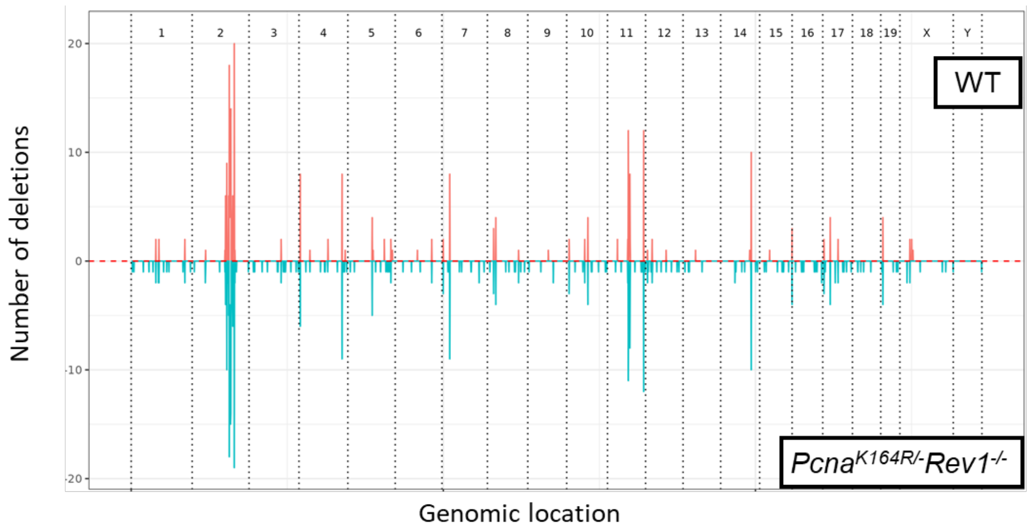


B

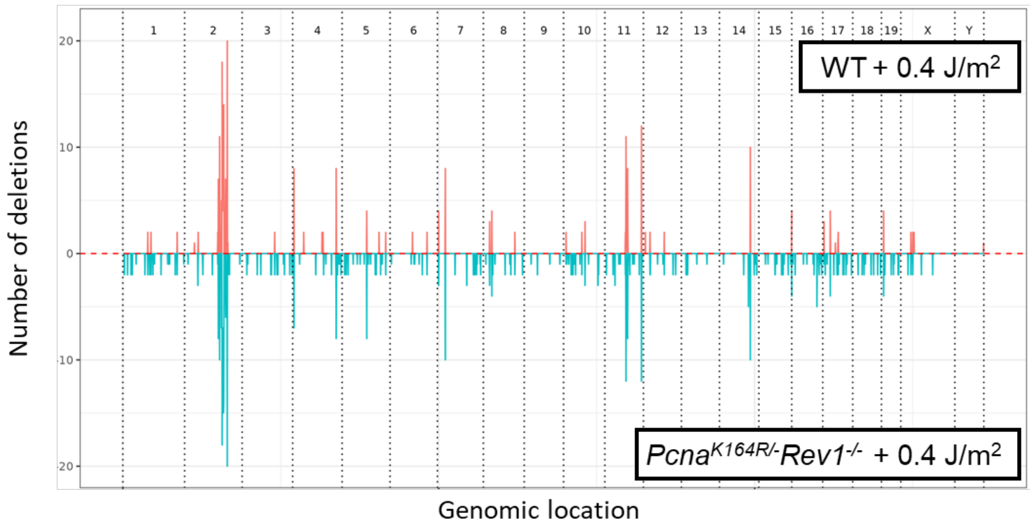


C

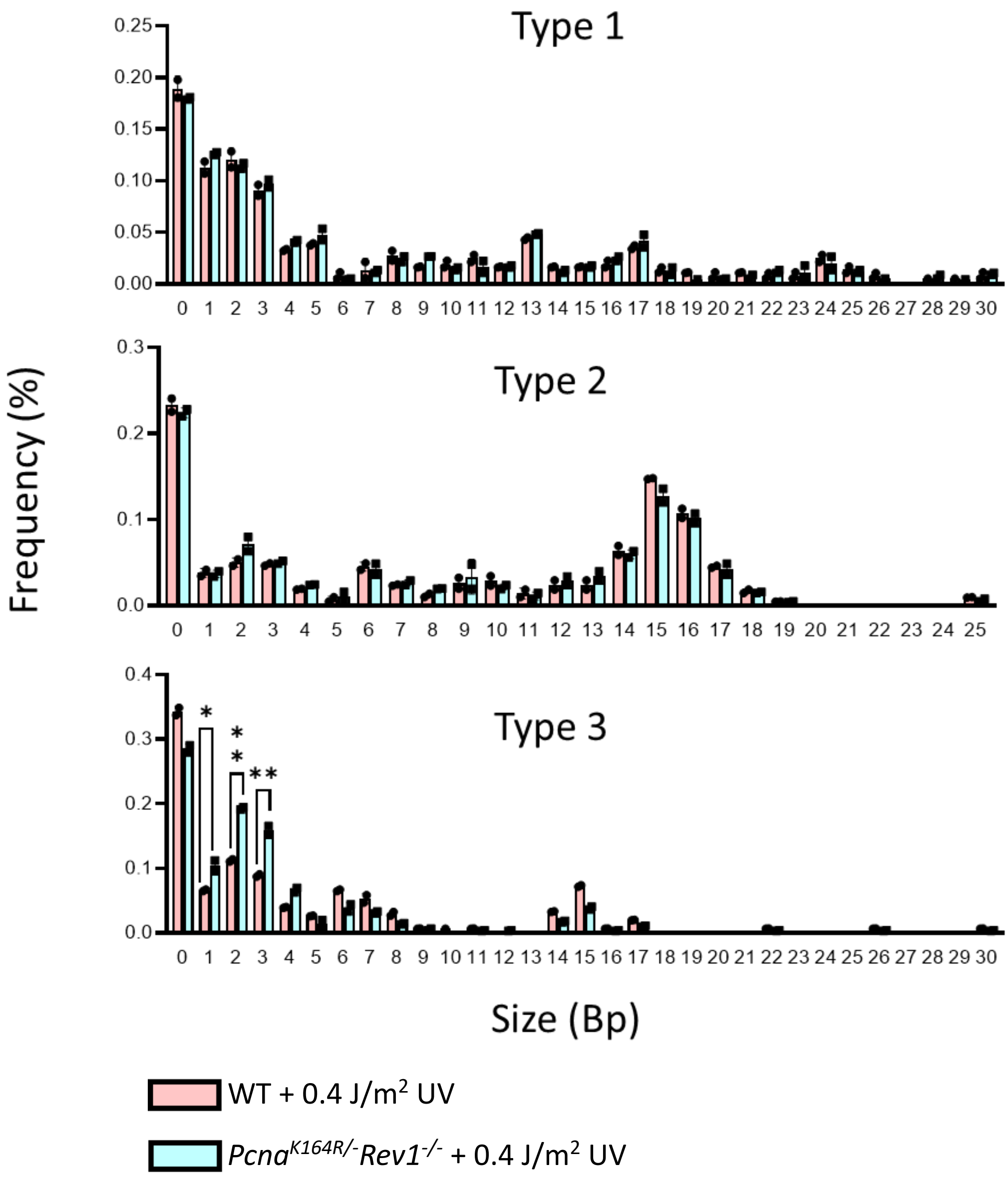

### Supplementary figure 6

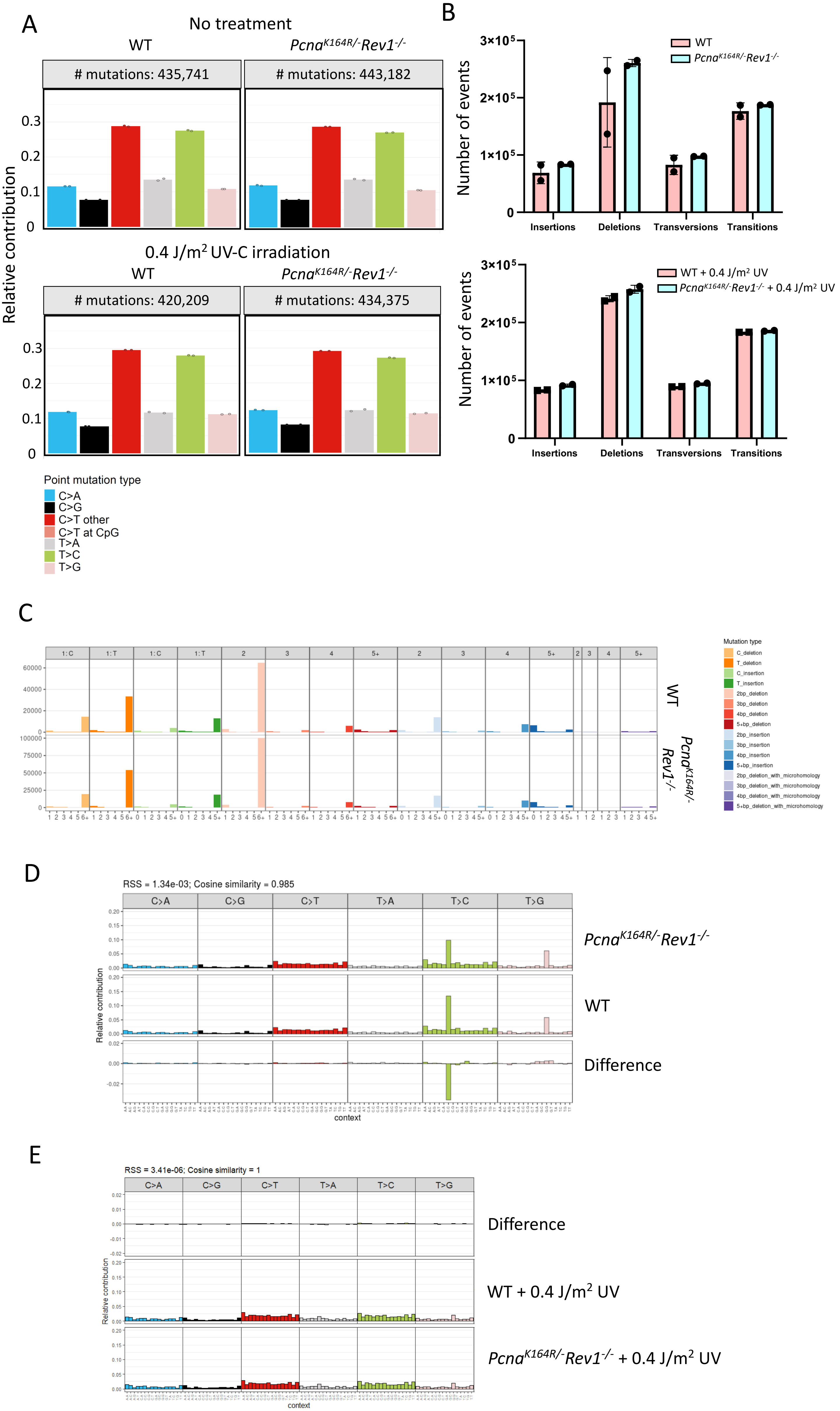

### Supplementary figure 7

A

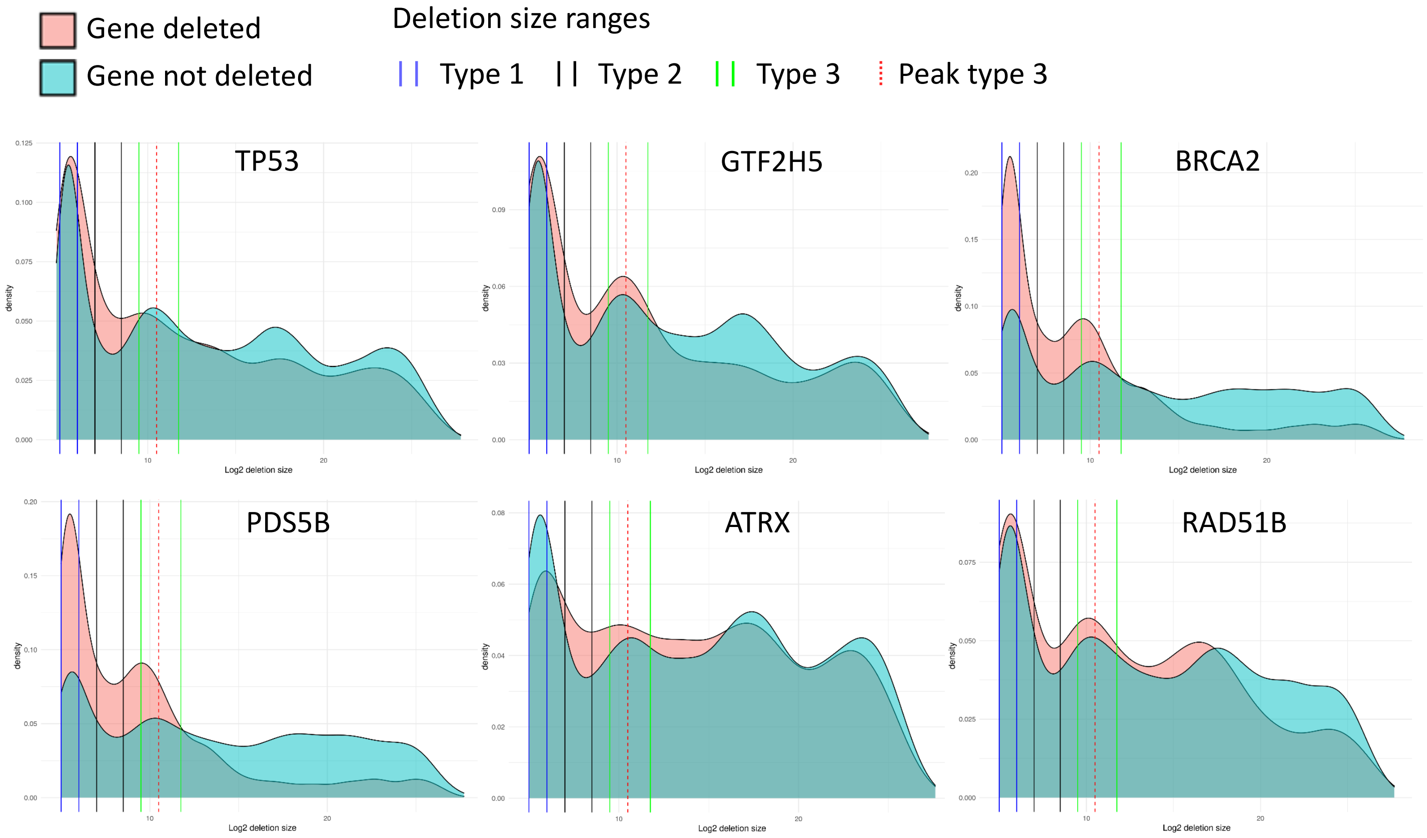


B

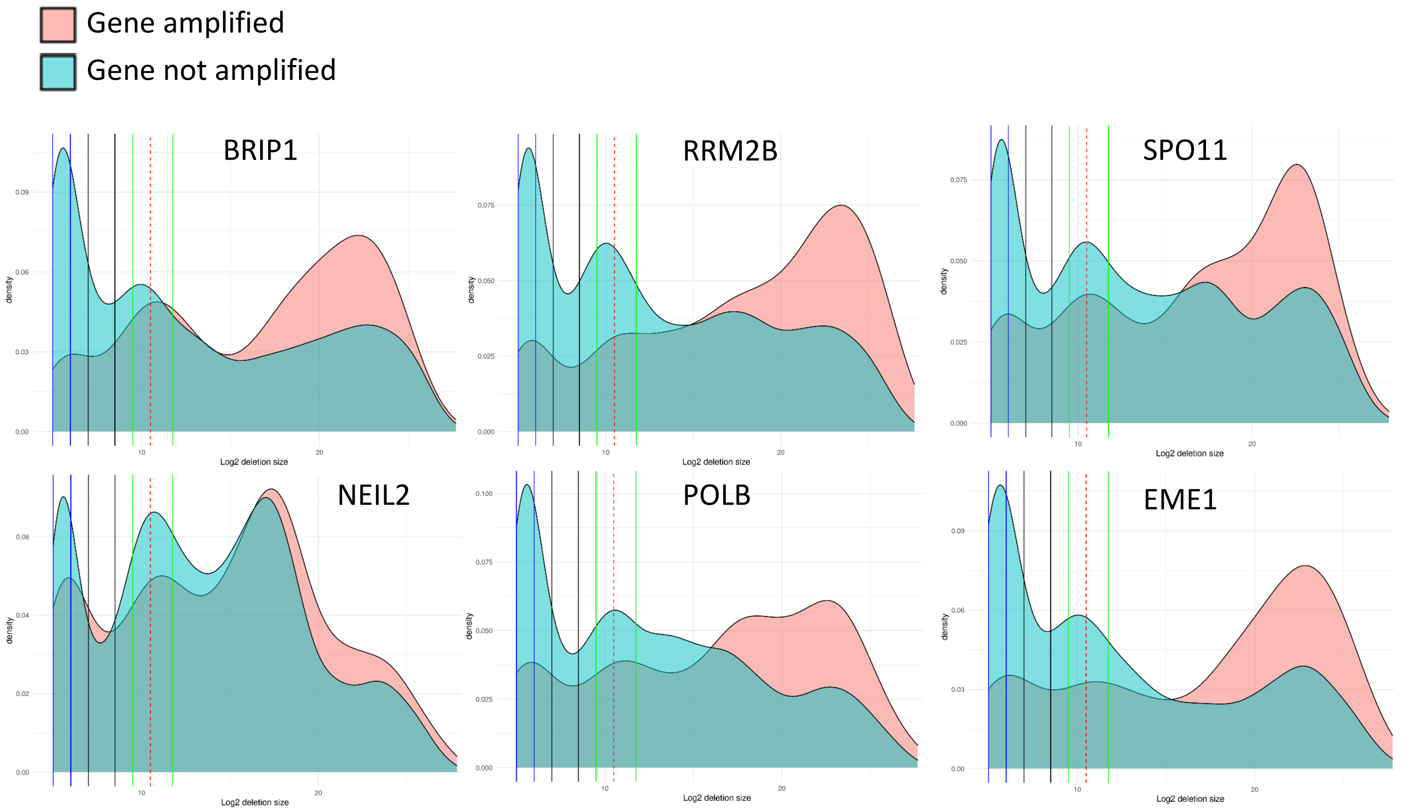

Supplementary figure 8

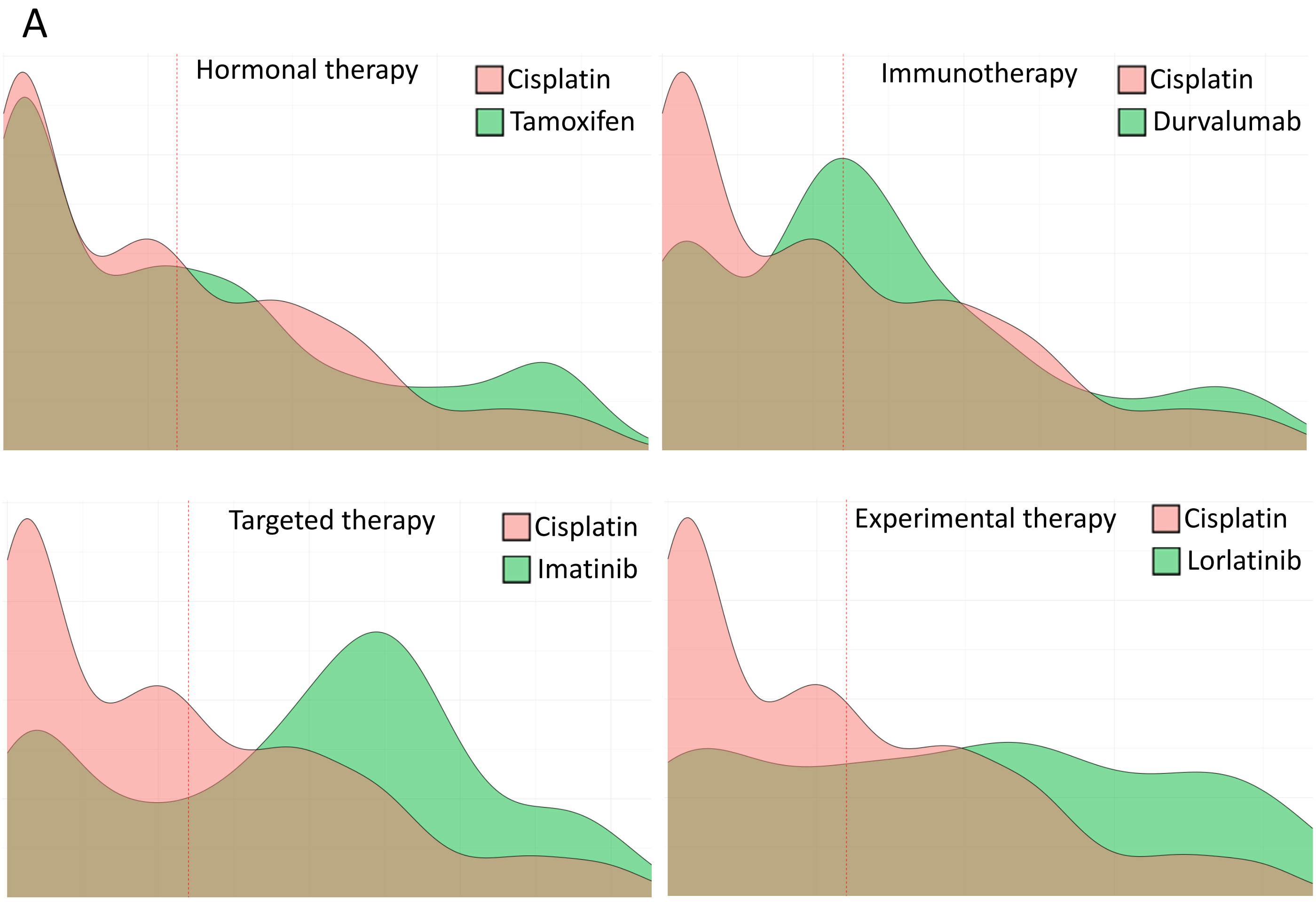

**Table S1: primers used for genotyping cells**

| **Oligonucleotides for PCRs and gRNAs** | **Sequence (5'-3')** | **Purpose** |
|---|---|---|
| **PCNA genotype FWD** | TGCAAGTGGAGAGCTTGGCAATG | Determine whether mice carry *Pcna*$^{K164}$ WT allele, or mutant *Pcna*$^{K164R}$ allele. |
| **PCNA genotype REV** | CTTTCCAAATGCTACCTGTggcg | Determine whether mice carry *Pcna*$^{K164}$ WT allele, or mutant *Pcna*$^{K164R}$ allele. |
| **REV1-KO genotype FWD** | GGCAACATGGCCAAGAAGAAC | Determine whether mice carry *Rev1* WT or KO allele. Identical for mice, MEFs, and lymphoma |
| **REV1-KO genotype REV** | TTATTCAGCTTGGCGAGCGCTTTTG | Determine whether mice carry *Rev1* WT or KO allele. Identical for mice, MEFs, and lymphoma |
| **REV1-KO genotype INT** | ACTCAGTCAGCAGACACATGC | Determine whether mice carry *Rev1* WT or KO allele. Identical for mice, MEFs, and lymphoma |

**Table S2: PCR settings for genotyping**

| **PCR settings** | **Time** |
|---|---|
| 95°C | 3min |
| 75°C | 5min |
| 72°C | 1.5min |
| Melting - 95°C (30-40 cycles) | 30sec |
| Annealing - 63°C (30-40 cycles) | 30sec |
| Extension - 72°C (30-40 cycles) | 45sec |
| Final extension - 72°C | 10min |
| Storage – 4°C-12°C | Until usage |
